# Bacterial cyclopropane fatty acid synthase mRNA is targeted by activating and repressing small RNAs

**DOI:** 10.1101/271551

**Authors:** Colleen M. Bianco, Kathrin S. Fröhlich, Carin K. Vanderpool

## Abstract

Altering membrane protein and lipid composition is an important strategy for maintaining membrane integrity during environmental stress. Many bacterial small RNAs (sRNAs) control membrane protein production, but sRNA-mediated regulation of membrane fatty acid composition is less understood. The sRNA RydC was previously shown to stabilize *cfa* (cyclopropane fatty acid synthase) mRNA, resulting in higher levels of cyclopropane fatty acids in the cell membrane. Here, we report that three additional sRNAs, ArrS, CpxQ, and GadF also regulate *cfa* post-transcriptionally. RydC, ArrS, and GadF act through masking an RNase E cleavage site in the *cfa* mRNA 5’ untranslated region (UTR), and all three sRNAs post-transcriptionally activate *cfa.* In contrast, CpxQ binds to a different site in the *cfa* mRNA 5’ UTR and represses *cfa.* Alteration of membrane lipid composition is a key mechanism for bacteria to survive low pH environments, and we show that *cfa* translation increases in an sRNA-dependent manner when cells are subjected to mild acid stress. This work suggests an important role for sRNAs in the acid stress response through regulation of *cfa* mRNA.

## Introduction

Bacteria modify the biophysical properties of their membranes to adapt to changing environmental conditions such as pH, temperature, and pressure fluctuations (Zhang & Rock, 2008). Membrane properties can be altered by changing the type or abundance of proteins embedded in the membrane or by modifying the relative proportions of different types of phospholipids. Membrane fluidity is a crucial biophysical property as it affects membrane-associated functions such as permeability to solutes, solute transport, and protein-protein interactions. The length and saturation of the acyl chains in the phospholipids determines the fluidity of the membrane, and bacteria adjust the ratio of saturated to unsaturated fatty acids (UFAs) to adapt to their environment. For example, bacteria can increase resistance to toxic compounds, such as antimicrobial peptides, by increasing the production of saturated fatty acids (Zhang & Rock, 2008), which will form a tightly packed and less fluid membrane compared to a membrane rich in unsaturated fatty acids.

While *de novo* production of fatty acids is an important adaptation mechanism, bacteria may encounter abrupt environmental changes that require rapid modification of the fatty acids already incorporated in the membrane. One post-synthetic modification is the conversion of a pre-existing UFA to a cyclopropane fatty acid (CFA) by the enzyme cyclopropane fatty acid synthase (encoded by *cfa).* CFAs are formed by the addition of a methylene group, derived from S-adenosyl methionine, across the double bond of an UFA incorporated in a phospholipid (Grogan & Cronan, 1997, Law, 1971). CFAs occur in the phospholipids of many species of bacteria, but although widely studied, their physiological function remains unclear. One hypothesis is that the formation of CFAs may reduce membrane fluidity and permeability (Grogan & Cronan, 1997), including the permeability to protons (Shabala & Ross, 2008). Indeed, *Escherichia coli cfa* mutants are unable to survive acid shock, a rapid shift from pH 7 to pH 3 (Brown, Ross et al., 1997, Chang & Cronan, 1999). In addition, pathogenic *E. coli* strains contain more CFAs and are more resistant to acid than nonpathogenic *E. coli* strains (Brown et al., 1997). The formation of CFAs has been further linked to bacterial virulence as inactivation of a *cfa* homologue that introduces a cyclopropane ring in the major cell envelope component (α-mycolates) of *Mycobacterium tuberculosis* leaves the bacteria unable to establish a persistent infection (Glickman, Cox et al., 2000).

Production of CFAs is regulated at multiple levels. In *E. coli* and *Salmonella, cfa* transcription is driven by two promoters (Kim, Kim et al., 2005, Wang & Cronan, 1994). The distal promoter is σ^70^-dependent and yields a long *cfa* transcript with a 212-nt 5’-untranslated region (UTR). The proximal promoter is controlled by the general stress response σ factor (σ^s^) encoded by *rpoS* and produces a short *cfa* transcript with a 34-nt 5’ UTR (Fig. 1A). The σ^70^-dependent promoter is functional throughout growth, whereas transcription from the σ^s^ promoter occurs only during stationary phase.

**Figure 1:**
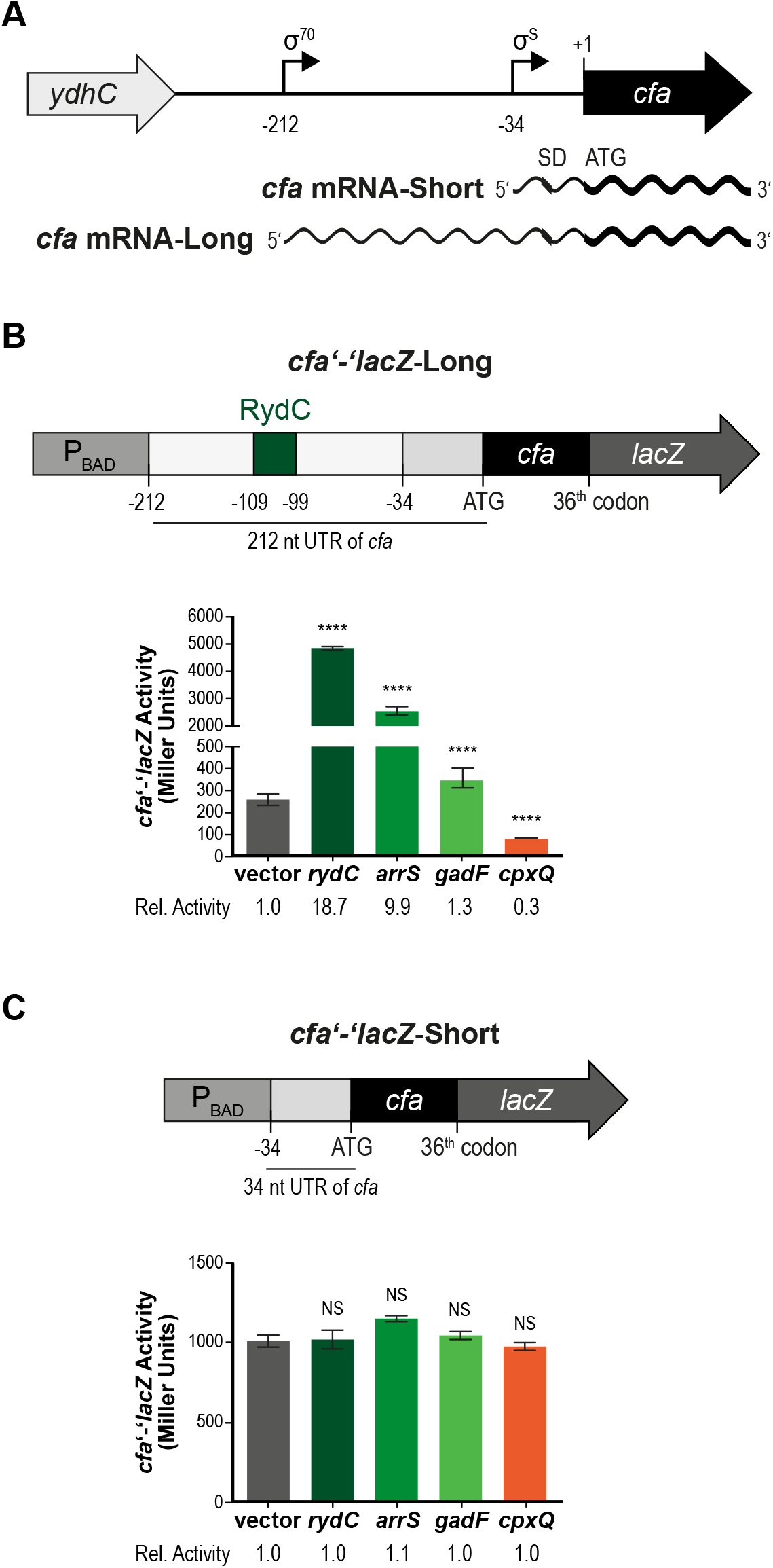
*cfa* expression is controlled by multiple sRNAs. (A) *cfa* has two promoters. Transcription from the distal promoter is σ^70^-dependent and yields a longer *cfa* transcript with a 212-nt 5’ UTR. Transcription from the proximal promoter is controlled by σ^s^ and produces a shorter *cfa* transcript with a 34-nt 5’ UTR. (B) A *cfa* translational fusion to *lacZ* (controlled by P_BAD_ promoter) was constructed. *cfa’-’lacZ-Long* fusion is from the distal σ^70^-dependent promoter which contains a 212-nt 5’ UTR that includes the RydC binding site (green square labeled “RydC”) and the sites predicted for the ArrS, CpxQ, and GadF. *cfa’-’lacZ*-Long carrying an empty or a P_lac_-*sRNA* plasmid were grown in TB medium with 0.002% L-arabinose to early exponential phase then sRNA expression was induced with 0.1 mM IPTG. Samples were harvested 60 minutes later and assayed for β-galactosidase activity of the reporter fusion. The error bars are standard deviations, and the statistical significance was determined using the Student’s *t*-test. \**P* < 0.05, \*\**P* < 0.005, \*\*\**P* < 0.0005, \*\*\*\**P* < 0.0001, ns is for not significant. The Student’s *t*-test was performed comparing fusion activity in response to expression of each sRNA to fusion activity in vector control. (C) A *cfa’-’lacZ*-Short fusion was constructed, which contains only proximal the σ^s^-dependent promoter and consequently not the predicted sRNA binding sites. Regulation of *cfa’-’lacZ*-Short by each sRNA was determined as described in 1B.

Post-transcriptional regulation by small non-coding RNAs (sRNAs) constitutes a major layer of gene expression control in bacteria. In many cases, a ubiquitous RNA chaperone, Hfq, is required for sRNA stability and to mediate base-pairing interactions between an sRNA and its cognate target mRNAs. A recent study applied a methodology based on ligation of Hfq-associated RNAs and sequencing of chimeric fragments to globally map RNA interactions *in vivo* (Melamed, Peer et al., 2016). This approach (termed RIL-seq) uncovered *cfa* mRNA interactions with multiple sRNAs in *E. coli:* ArrS, CpxQ, GadF, and RydC. RydC was previously reported to stabilize *cfa* mRNA, which in turn increases levels of cyclopropane fatty acid (CFA) synthase and results in proportionally more CFAs in membrane lipids (Fröhlich, Papenfort et al., 2013). RydC, a 64 nucleotides (nt) long sRNA, folds as a characteristic pseudoknot structure which exposes a stretch of highly conserved nucleotides at the very 5’ end of the RNA. Binding of RydC within the long *cfa* mRNA masks a recognition site for the major endonuclease RNase E and inhibits transcript degradation. In addition, RydC has been reported to regulate *yejABEF* (Antal, Bordeau et al., 2005), *csgD* (Bordeau & Felden, 2014), *pheA* and *trpE* (King, Vanderpool et al., 2019) mRNAs, though the conditions and signals stimulating RydC production and the physiological role of RydC in bacterial physiology remain unknown. In contrast, ArrS, CpxQ and GadF sRNAs were previously linked to cell envelope or acid stress responses. Production of the CpxQ sRNA depends on the Cpx two-component system that responds to cell envelope stress (Chao & Vogel, 2016). CpxQ is processed by RNase E from the 3’ UTR of *cpxP* mRNA, which encodes a chaperone of the Cpx system (Chao & Vogel, 2016). Together with Hfq, CpxQ represses translation of multiple targets many of which encode inner membrane proteins (Chao & Vogel, 2016, Grabowicz, Koren et al., 2016). ArrS is an antisense sRNA that is encoded upstream of *gadE* (encoding the major acid resistance transcription factor) and is complementary to the 5’ UTR of the longest of three *gadE* transcripts (Aiso, Murata et al., 2011). ArrS expression is induced by low pH through σ^s^ and GadE, and overexpression of ArrS increases survival of cells exposed to acidic pH (Aiso, Kamiya et al., 2014). A second sRNA associated with the *gadE* locus is GadF, which is liberated from the 3’ UTR of *gadE* mRNA by endonucleolytic cleavage (Melamed et al., 2016). To date, GadF has not been extensively studied, however the enrichment of acid response-associated genes in its RIL-seq profile suggests a role in this pathway.

In this study, we sought to elucidate the mechanisms of regulation of *cfa* mRNA by the four sRNAs, RydC, ArrS, GadF, and CpxQ, and to gain insight into the physiological importance of sRNA-mediated regulation of membrane CFA composition in the model organism *E. coli.* Our results indicate that RydC, ArrS and GadF activate *cfa* to varying degrees, and all three require overlapping binding sites in the long 5’ UTR of *cfa* mRNA. We found that RydC is responsible for increasing *cfa* expression at acidic pH, and that cells exhibit an acid stress resistance phenotype that is dependent on sRNA-mediated post-transcriptional activation of *cfa.* Unlike RydC, ArrS and GadF, the sRNA CpxQ represses *cfa* post-transcriptionally. CpxQ binds at a distinct site on *cfa* mRNA, upstream from the site bound by activating sRNAs. Our results suggest that CpxQ binding at this site modulates the stability of *cfa* mRNA by promoting its degradation by RNase E. Altogether, our work suggests that several distinct sRNAs, presumably responding to different environmental conditions, can modulate membrane fatty acid composition via activation or repression of *cfa* mRNA.

## Results

### Multiple sRNAs regulate CFA synthase mRNA in an isoform-specific manner

At the transcriptional level, expression of *cfa* is controlled by the activity of two promoters that yield mRNAs with a 212-nt UTR (distal σ^70^-dependent promoter) or a 34-nt UTR (σ^S^-dependent promoter), respectively (Wang & Cronan, 1994) (Fig. 1A). Only the long mRNA isoform is subject to post-transcriptional regulation by the sRNA RydC, which stabilizes the transcript by base-pairing at an RNase E recognition site within the 5’ UTR (Fröhlich et al., 2013). An experimental study based on the ligation of Hfq-associated RNAs (Melamed et al., 2016) confirmed the previously observed interaction between *cfa* mRNA and RydC sRNA, and in addition revealed the potential interactions of the *cfa* transcript to ArrS, CpxQ, and GadF sRNAs. To verify that ArrS, CpxQ, and GadF sRNAs alter *cfa* expression post-transcriptionally *in vivo,* we constructed two translational *cfa’-’lacZ* fusions under the control of the arabinose-inducible P_BAD_ promoter (King et al., 2019). One fusion contains the 212-nt 5’ UTR and the first 36 codons of *cfa (cfa’-’lacZ-Long,* Fig. 1B), while the second has only the short 34-nt 5’ UTR *(cfa’-’lacZ-* Short, Fig. 1C). We ectopically induced each sRNA from a plasmid (P_Lac_ promoter), and determined the effect on the two reporter fusions. RydC and ArrS increased *cfa’-’lacZ-*Long activity 18-and 10-fold, respectively, while CpxQ repressed the long fusion ~3-fold (Fig. 1B). Expression of GadF resulted in a very minor increase (1.3-fold) of *cfa’-’lacZ-*Long activity (Fig. 1B). No sRNA had an effect on *cfa’-’lacZ*-Short activity (Fig. 1C). These results indicate that the long isoform of *cfa* mRNA can be post-transcriptionally activated or repressed by multiple sRNAs, while the short isoform is not affected by these regulatory molecules.

Furthermore, RIL-seq also revealed the potential interactions of the *cfa* transcript to OxyS and GcvB sRNAs. We tested *cfa’-’lacZ*-Long and *cfa’-’lacZ*-Short for regulation by OxyS and GcvB (Fig. EV1). Expression of OxyS increased *cfa’-’lacZ*-Long activity 2.1-fold while expression of GcvB decreased *cfa’-’lacZ*-Long activity 1.5-fold (Fig. EV1A). Neither sRNA significantly affected *cfa’-’lacZ*-Short activity (Fig. EV1B). OxyS is expressed under oxidative stress and represses *rpoS* expression by an unknown mechanism (Altuvia, WeinsteinFischer et al., 1997, Zhang, Altuvia et al., 1998). GcvB mutants are acid sensitive and this sensitivity may be due to reduced RpoS levels, however how GcvB activates RpoS expression is not known (Jin, Watt et al., 2009). Due to the potential regulation of RpoS by OxyS and GcvB (which could indirectly regulate *cfa’-’lacZ*-Long by modulating expression from the σ^S^-dependent promoter), and the observed minor effect of these sRNAs on regulation of *cfa,* we did not further characterize regulation by OxyS and GcvB.

### Activating and repressing sRNAs bind distinct sites on *cfa* mRNA

RydC, ArrS, GadF, and CpxQ all regulated the *cfa* translational fusion containing the *cfa* 212-nt 5’-UTR but not the *cfa* translational fusion containing the short 34-nt 5’-UTR. Given that sRNAs typically regulate their targets by engaging in Hfq-dependent formation of base-pairing interactions, we assumed that each sRNA recognizes a site within the upstream region of *cfa* mRNA. RydC has previously been shown to base-pair at an RNase E recognition site located within this region (Fröhlich et al., 2013). We used IntaRNA (Mann, Wright et al., 2017) to predict interactions between ArrS, GadF, and CpxQ with *cfa* mRNA, and determined that all three sRNAs potentially bound the target in proximity to the previously determined RydC site (Figs. 2A, EV2A). To genetically test these base pairing interactions we first constructed a *cfa’-’lacZ* fusion with a 133-nt deletion from the beginning of the 212-5’ UTR, which contains predicted binding sites of all of the sRNAs (cfa’-’lacZ-ΔsRNABS, Fig. EV2B), including the RNase E recognition site that RydC protects. This fusion only contains the proximal σ^S^-dependent promoter and 79-nt upstream of this promoter. Of note, *cfa’-’lacZ*-ΔsRNABS had higher basal activity compared to *cfa’-’lacZ*-Long (Figs. 1B, EV2C), presumably because *cfa’-’lacZ-* ΔsRNABS fusion does not contain a known RNase E recognition site, inherently stabilizing the transcript. When each individual sRNA was ectopically expressed from a plasmid, none of the sRNAs affected *cfa’-’lacZ*-ΔsRNABS activity (Fig. EV2C), indicating each sRNA base pairs upstream of the −79 nt position. To specifically analyze where each sRNA binds *cfa* mRNA, we completed structure probing experiments (Fig. 2). We digested a ~140-nt fragment of *cfa* mRNA (−212 to −72 relative to the start codon) in the presence of Hfq and each of the four individual candidate sRNAs with ribonuclease T1 or lead (II) acetate (Fig. 2B). We found that RydC (lanes 5, 10), and ArrS (lanes 6, 11) bind sites that overlap, whereas CpxQ (lanes 8,13) binds upstream at a distinct site. Protection by GadF was weak (lane 7, T1 digest) or not observed (lane 12, lead (II) acetate cleavage), consistent with *in vivo* observations that GadF is a very weak regulator of *cfa* mRNA. We did not characterize GadF-cfa mRNA interactions further. The results of the *in vitro* structure probing experiment were in line with our computational predictions of base-pairing sites, suggesting that the two activating sRNAs, ArrS and RydC, shared a common binding site whereas CpxQ employed a distinct site to repress *cfa* mRNA.

**Figure 2:**
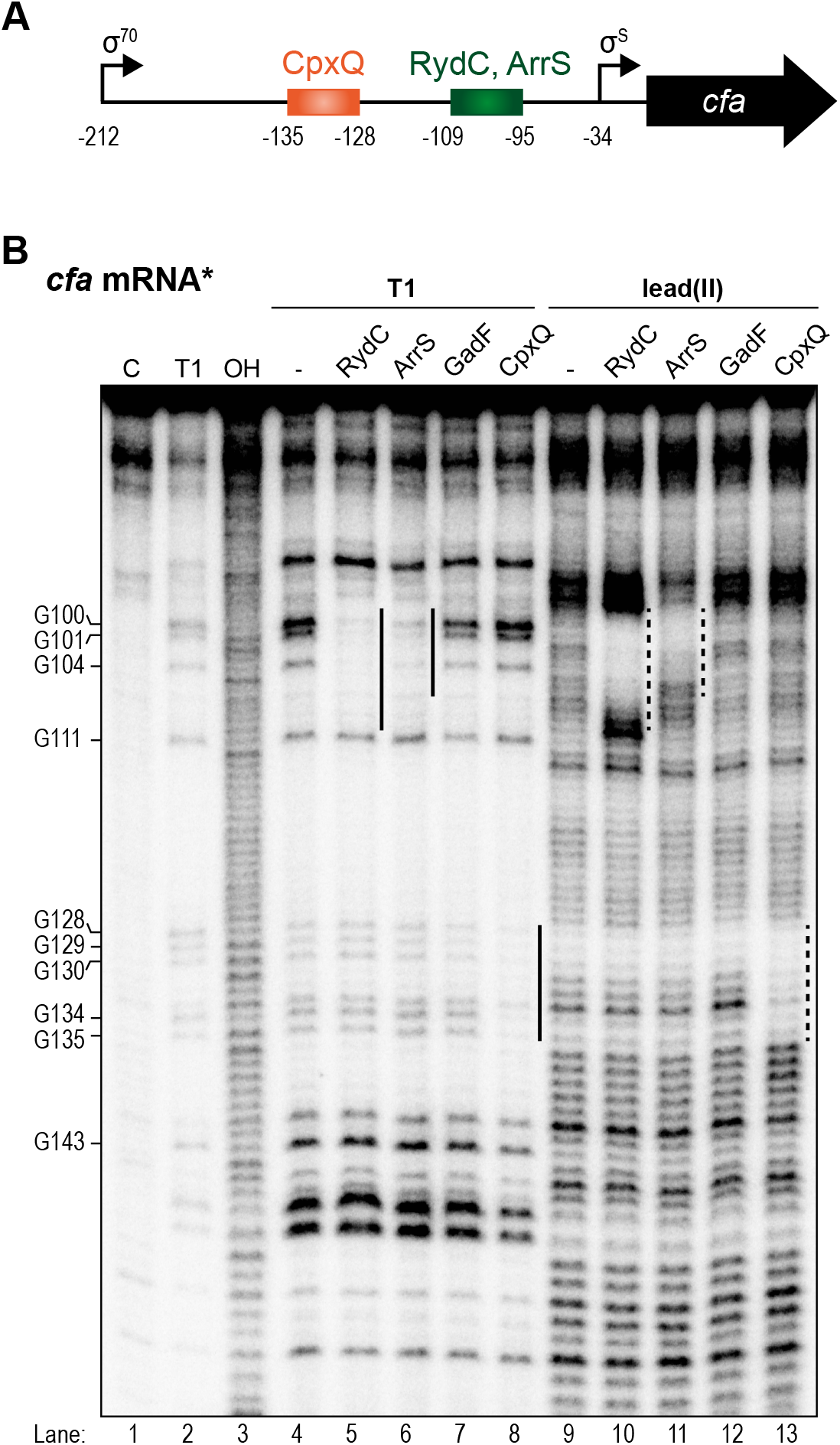
Activating and repressing sRNAs bind different sites on *cfa* mRNA. (A) Predicted base pairing sites of sRNAs on *cfa* mRNA. (B) *In vitro* structure probing using 5’ end-labeled *cfa* mRNA with RNase T1 (lanes 4-8) and lead(II) acetate (lanes 9-13) in the presence Hfq (20 nM) and each sRNA (200 nM). RNase T1 and alkaline ladders of cfa mRNA were used to map cleaved fragments. Positions of G-residues are indicated relative to the translational start site. Each sRNA binding site is marked with a black line right of the lane.

To test base-pairing predictions with each sRNA directly, we did a mutational analysis. We first introduced a point mutation in the two activating sRNAs, ArrS and RydC, that were predicted to disrupt base pairing to *cfa’-’lacZ*-Long (C7G for *arrS* M1), and C5G for *rydC* M1 (Fig. 3A). These mutated sRNA variants, ArrS M1 and RydC M1, could no longer activate *cfa’-’lacZ*-Long (Fig. 3B). We made a compensatory mutation (G101C) to create *cfa’-’lacZ*-Long M1, to restore putative base pairing interactions with the mutant sRNAs (Fig. 3A, B). Wild-type ArrS and RydC could not activate the mutant *cfa’-’lacZ*-Long M1fusion (Fig. 3B), consistent with the idea that this residue in the *cfa* 5’-UTR is critical for interactions with both sRNAs. In contrast, the mutants RydC M1 and ArrS M1 activated the *cfa’-’lacZ*-Long M1 fusion (Fig. 3B), with fold-activation restored to levels similar to the wild-type mRNA-sRNA pairs (Fig. 3B).

**Figure 3:**
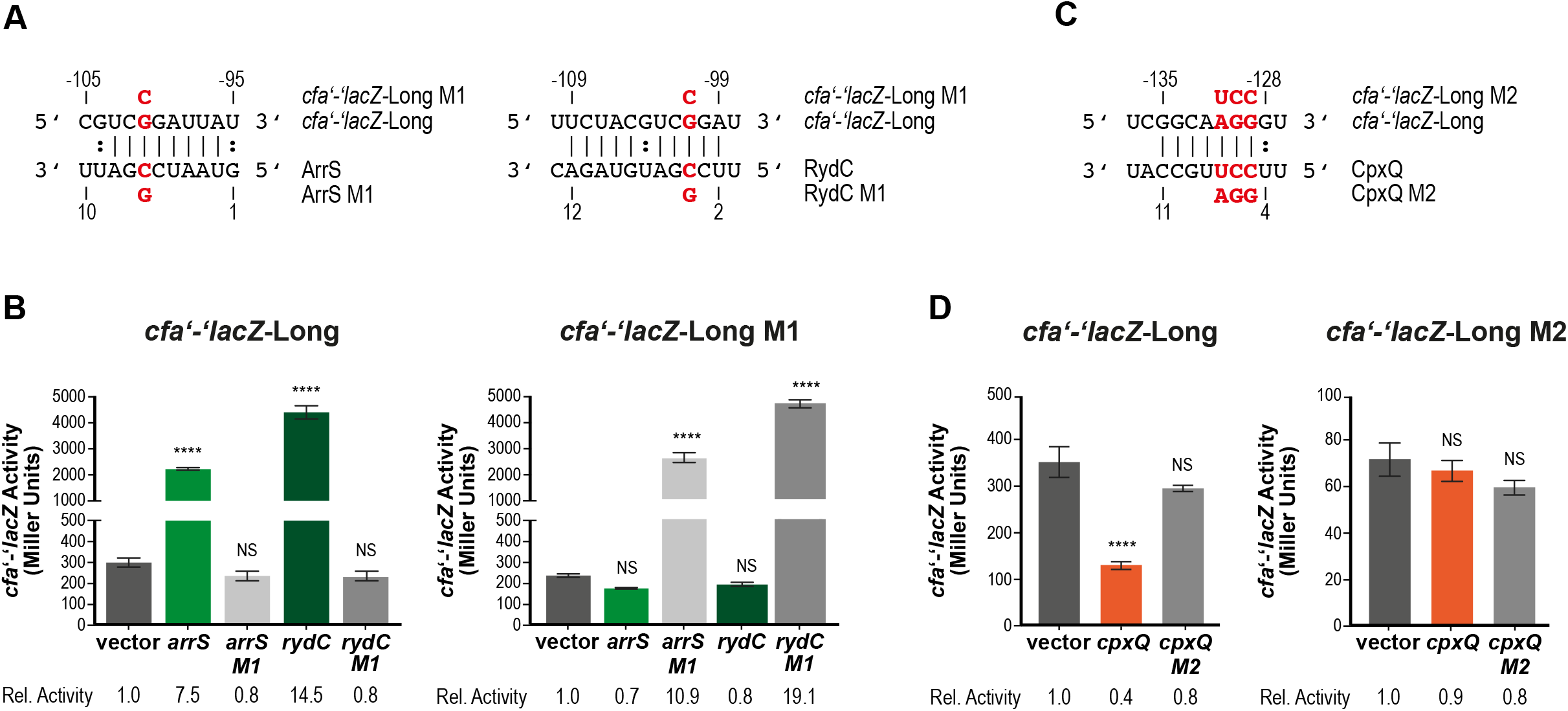
Activating and repressing sRNAs bind different sites on *cfa* mRNA. (A) Predicted base pairing between *cfa* mRNA and ArrS or RydC. The red nucleotides were mutated to test base pairing. (B) Left: Mutant sRNAs (ArrS M1, RydC M1) and WT sRNAs were tested for activation of *cfa’-’lacZ*-Long as described in Fig. 1B. (B) Right: One point mutation (G101C) were made in *cfa’-’lacZ*-Long (called *cfa’-’lacZ-Long* M1). Mutant sRNAs (ArrS M1, RydC M1) and WT sRNAs were tested for activation of *cfa’-’lacZ*-Long M1 as described in Fig. 1B. (C) Base pairing between *cfa* mRNA and CpxQ. The red nucleotides were mutated to test base pairing. (D) Left: Mutant CpxQ M2 and WT CpxQ were tested for activation of *cfa’-’lacZ*-Long as described in Fig. 1B. (D) Right: Three point mutations (A131T, G130C, G129C) were made in *cfa’-’lacZ*-Long (called *cfa’-’lacZ*-Long M2). Mutant CpxQ M2 and WT CpxQ were tested for activation of *cfa’-’lacZ*-Long M2 as described in Fig. 1B.

To characterize CpxQ-mediated regulation of *cfa* mRNA, we mutated the predicted CpxQ binding site (Fig. 2A, 3C). We introduced three point mutations in CpxQ that should disrupt base pairing to *cfa’-’lacZ*-Long (C5G, C6G, U7A, called CpxQ M2, Fig. 3C). This CpxQ mutant lost most of the repression activity for the *cfa’-’lacZ*-Long (Fig. 3D). Mutations in the *cfa* fusion (U103A, C102G, G101C, called *cfa’-’lacZ*-Long M2) that disrupt base pairing interactions with wild-type CpxQ, likewise eliminated the repression by wild-type CpxQ (Fig. 3D). We noted that the point mutations in the *cfa* fusion that disrupt the interaction with CpxQ strongly reduce the basal levels of reporter activity (~5-fold). This suggests that these nucleotides are important for stability of *cfa* mRNA. Wild-type levels of repression were not fully restored in the compensatory mutant pair – CpxQ M2 and *cfa’-’lacZ*-Long M2 (Fig. 3D), suggesting that the mutation in the fusion already serves to make *cfa* mRNA more sensitive to degradation such that CpxQ binding has no further impact. Nevertheless, the loss of regulation caused by individual mutations (Fig. 3D) and the clear CpxQ footprint on *cfa* mRNA (Fig. 2B) strongly support that CpxQ binds at a site on *cfa* mRNA substantially upstream of the binding site of the activating sRNAs.

### Activating and repressing sRNAs modulate RNase E-dependent degradation of *cfa* mRNA

Previous studies determined that the RydC-cfa mRNA base pairing prevents RNase E-mediated decay and stabilizes the mRNA to allow increased translation (Fröhlich et al., 2013). We hypothesize, because ArrS pairs with the same region of *cfa* mRNA as RydC, that ArrS regulates *cfa* mRNA stability by the same mechanism. However, the repressive effect of CpxQ must be based on a different mechanism. CpxQ has been shown to repress other mRNA targets using one of two conserved seed regions (Chao & Vogel, 2016). One of these seed regions matches the region of CpxQ that base pairs with *cfa* mRNA (Fig. 3C). CpxQ represses its other targets by base pairing near the Shine-Dalgarno sequence to prevent ribosome binding and directly inhibit translation initiation, or by base pairing within the coding region and stimulating mRNA decay by RNase E (Chao & Vogel, 2016). Since the CpxQ-cfa interaction site within the *cfa* 5’-UTR is far upstream of the Shine-Dalgarno sequence CpxQ must not repress translation initiation directly. Instead, we wanted to determine if CpxQ caused *cfa* mRNA decay by stimulating RNase E activity. When CpxQ is produced from the native chromosomal locus, it carries a 5’ monophosphate (5’P) end because it is a processed product of the *cpxP* mRNA and not a primary transcript, which would carry a 5’ triphosphate (5’PPP). It has been shown that the 5’P end of an sRNA can stimulate RNase E activity and lead to rapid degradation of the paired mRNA (Bandyra, Said et al., 2012). It was demonstrated *in vitro* that RNase E-dependent degradation of another CpxQ target, *nhaB* mRNA, was faster in the presence of 5’P CpxQ compared to 5’PPP CpxQ (Chao & Vogel, 2016). To determine whether the phosphorylation status of the CpxQ 5’ end impacts its ability to regulate *cfa* mRNA, we designed CpxQ plasmids that would allow for the production of processed 5’P CpxQ. Three plasmids were used (Fig. EV3A). All three plasmids contained DNA regions encompassing mature CpxQ and its ρ-independent terminator. The transcription start site for each plasmid varied (Fig. EV3A). The CpxQ plasmid (the same as used in experiments described above) placed the +1 of CpxQ at the native processing site. The CpxQ1 plasmid contained a larger region, beginning in the *cpxP* coding region (just after the *cpxP* start codon). The CpxQ2 plasmid contained a region starting 30-nt upstream of the processing site in the 3’ UTR of *cpxP.* We expected the CpxQ plasmid would produce CpxQ containing a 5’PPP while the CpxQ1 and CpxQ2 plasmids would produce a *cpxQ* transcript that would be processed by RNase E and yield CpxQ with a 5’P. We tested how each plasmid regulated *cfa’-’lacZ*-Long (Fig. EV3B) and found that both CpxQ1 and CpxQ2 repressed *cfa’-’lacZ*-Long activity to the same extent as CpxQ. A northern blot probing for CpxQ confirmed that CpxQ1 and CpxQ2 were processed to produce the 58nt CpxQ and that all three plasmids produced the same amount of the 58nt CpxQ sRNA (Fig. EV3C). In addition, deletion of *rppH* (encoding the 5’ pyrophosphatase which converts 5’ PPP-CpxQ to 5’ P-CpxQ) in the *cfa’-’lacZ*-Long fusion background did not affect regulation by CpxQ (Fig. EV3D). These results suggest that the status of the CpxQ 5’ end is not critical for its regulatory activity on *cfa* mRNA.

We anticipated that CpxQ modulates the stability of *cfa* mRNA by making it more susceptible to cleavage by RNase E. The RNase E cleavage site protected by RydC is far downstream of the CpxQ binding site. Given the AU-rich nature of the *cfa* 5’ UTR, we hypothesized that there might be additional RNase E cleavage sites that are normally hidden in secondary structure but are made available to be recognized by RNase E upon CpxQ binding. We completed a primer extension experiment to profile the mRNA cleavage products of the *cfa* 5’ UTR in response to RydC and CpxQ (Fig. 4A). The two *cfa* transcription start sites are the most dominant signals. RydC strongly activates the longer cfa1 mRNA but has no effect on the shorter *cfa2* transcript (lanes 7-9). Furthermore, the mRNA cleavage product that corresponds to the RydC-cfa interaction site disappears when RydC is expressed (compare lanes 7 and 9), agreeing with previous work that this area contains an RNase E cleavage site that is masked upon formation of the RydC-cfa RNA duplex (Fröhlich et al., 2013). CpxQ repressed the longer cfa1 mRNA and had no effect on the shorter *cfa2* transcript (lanes 10-12). No other differences in band signals were detected, indicating that we were unable to detect new CpxQ-dependent mRNA cleavage products.

**Figure 4:**
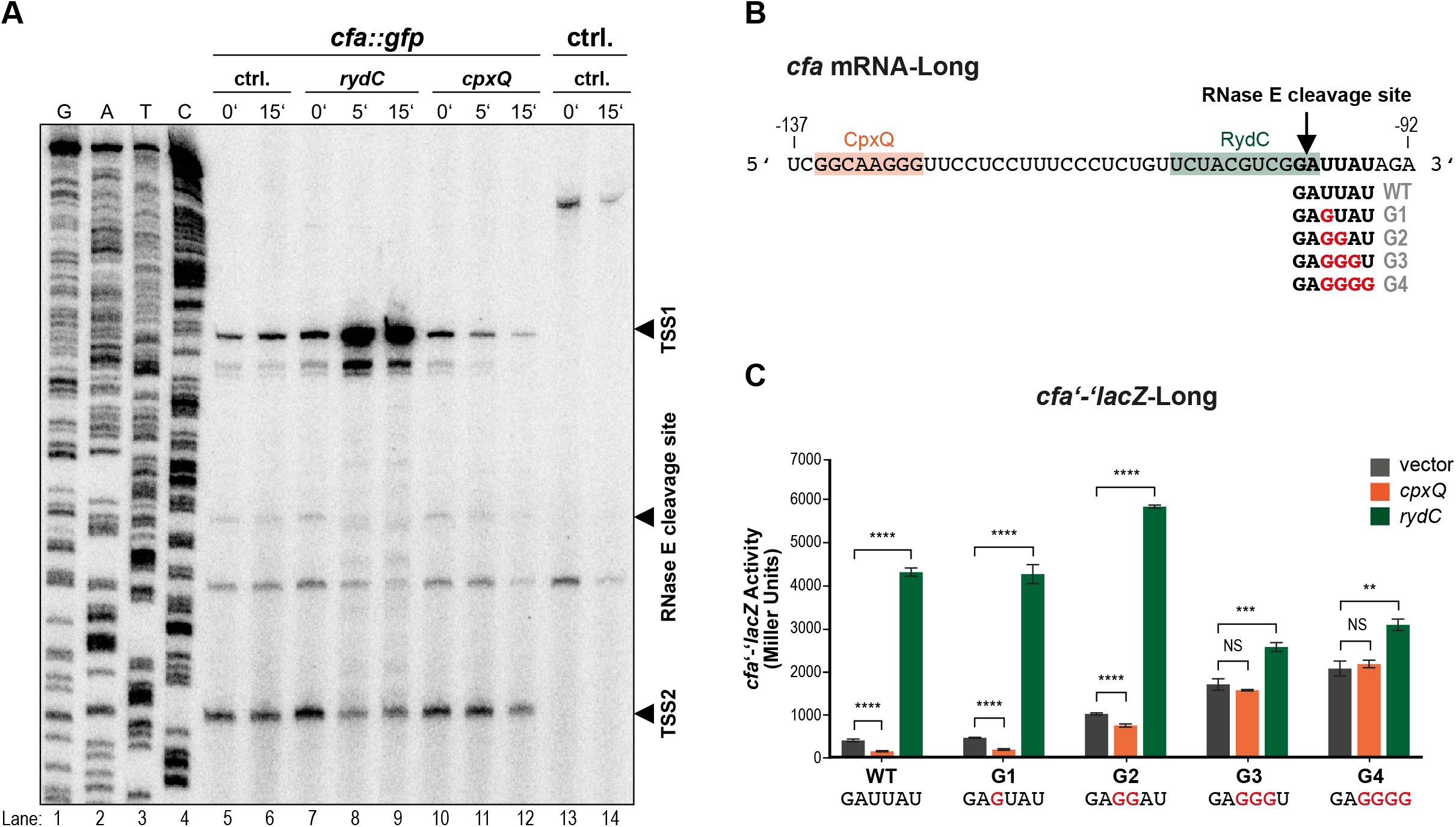
Activating and repressing sRNAs modulate RNase E-dependent degradation of *cfa* mRNA. (A) Primer extension of 5’ UTR of *cfa* mRNA reveals no CpxQ-dependent mRNA cleavage products. RNA samples were withdrawn prior to and 5 or 15 minutes after arabinose addition to induce vector control (lanes 5-6), RydC (lanes 7-9), or CpxQ (lanes 10-12). RNA samples were used as templates for primer extension using a 5’ end-labeled *cfa* primer. Transcripts were identified using cfa-specific sequencing ladders (lanes 1-4). Transcription start site 1 (TSS1) and 2 (TSS2) and the known RNase E cleavage site are marked with solid arrows. B) Multiple mutations of the RNase E recognition site on *cfa* mRNA were made in the *cfa’-’lac*Z-Long fusion background (called G1-G4). Black arrow indicates RNase E cleavage site. CpxQ and RydC base pairing locations are indicated. (C) Regulation of *cfa’-’lacZ*-Long (WT) or *cfa’-’lacZ*-Long G1-G4 by CpxQ and RydC was tested as described in Fig. 1B.

Another possibility is that CpxQ modulates the stability of *cfa* mRNA by making it more susceptible to cleavage at the known RNase E recognition site that is protected by RydC. To test whether RNase E-dependent cleavage is involved in CpxQ-mediated repression of *cfa,* we made mutations in putative RNase E sites in the *E. coli cfa* 5’ UTR (Fig. 4B). A global study of RNase E cleavage sites predicted a 5-nt minimal RNase E consensus sequence of “RNs↓WUU” (with R as G/A, W as A/U, N as any nucleotide and ↓ as the cleavage site) (Chao, Li et al., 2017). Moreover, in *Salmonella,* mutation of a U residue 2-nt downstream of the mapped RNase E cleavage site (Fig. EV4) significantly reduced the cleavage of *cfa* by RNase E (Chao et al., 2017). This uridine is conserved in *E. coli cfa* mRNA, so we mutated this U to a G (a non-preferred nucleotide for RNase E recognition) in the *cfa’-’lacZ*-Long (“G1” mutant, Fig. 4B, C,) and tested for regulation by RydC and CpxQ. Mutation of the U alone in the G1 mutant did not reduce the basal level of the fusion compared to WT, suggesting that in *E. coli,* the mutation did not substantially alter RNase E cleavage of *cfa* mRNA (Fig. 4C). CpxQ and RydC were able to regulate the G1 mutant similar to the wild-type fusion (Fig. 4C).

In some cases, uridines at the second and third position downstream of an RNase E cleavage site are highly conserved and mutation of both U residues is required to fully inhibit cleavage by RNase E (Chao et al., 2017). The sequence surrounding the RNase E cleavage site in the 5’ UTR of *cfa* differs between *Salmonella* and *E. coli* (Fig. EV4). In *Salmonella,* the sequence surrounding the RNase E cleavage site is GG↓AUA while in *E. coli* it is GG↓AUU. Changing both U residues to Gs (the G2 mutant, Fig. 4B, C) increased the basal activity of *cfa’-’lacZ*-Long (~2-fold compared to WT) suggesting that these mutations inhibit cleavage by RNase E (Fig. 4C). The G2 mutation did not impair activation by RydC, but substantially impaired CpxQ-mediated repression (Fig. 4C). Mutation of the next two residues downstream of the RNase E cleavage site yielded “G3” and “G4” mutant fusions (Fig. 4B). The basal levels of activity were further increased in these fusions, and both activation by RydC and repression by CpxQ were abrogated (Fig. 4C). These results strongly suggest that the regulation of *cfa* mRNA by both activating and repressing sRNAs involves modulation of accessibility of a single RNase E cleavage site that is adjacent to the site for base pairing by activating sRNAs, but far from the binding site of the repressing sRNA.

### Role of Hfq in sRNA-dependent regulation of *cfa* mRNA

In the previous study of RydC-mediated regulation of *cfa* mRNA, it was shown that swapping the RydC binding site on *cfa* mRNA for the binding site of another Hfq-dependent sRNA (e.g., RybB or RyhB) reprogrammed the *cfa* mRNA to be activated by the corresponding sRNA (Fröhlich et al., 2013). However, swapping the RydC binding site with an Hfq-independent sRNA binding site (i.e., IsrA) only had a small activating effect (~two-fold vs. ten-fold by RydC) (Fröhlich et al., 2013). This observation suggested that Hfq plays a key role in sRNA-dependent regulation of *cfa* mRNA stability. An Hfq binding site at the very 5’ end of the UTR of *Salmonella cfa* mRNA (Fig. 5A) was identified by CLIP-seq (Holmqvist, Wright et al., 2016). To further characterize the effects of this Hfq binding site on sRNA-dependent regulation of *cfa* mRNA, we deleted the first 50-nt of the 5’-UTR (−212 to −163, Fig. 5A, B, called *cfa’-’lacZ-ΔHfqBS)* which includes the Hfq binding site. This fusion had lower basal activity compared to *cfa’-’lacZ-*Long (~2-fold decrease; Fig. 5B), suggesting that the Hfq binding site located in the −212 to −163 region of the *cfa* mRNA plays a role in *cfa* mRNA structure or stability. Regulation of *cfa’-’lacZ*-ΔHfqBS by both activating and repressing sRNAs was strongly impaired even though the fusion retains both sRNA binding sites (Fig. 5B).

**Figure 5:**
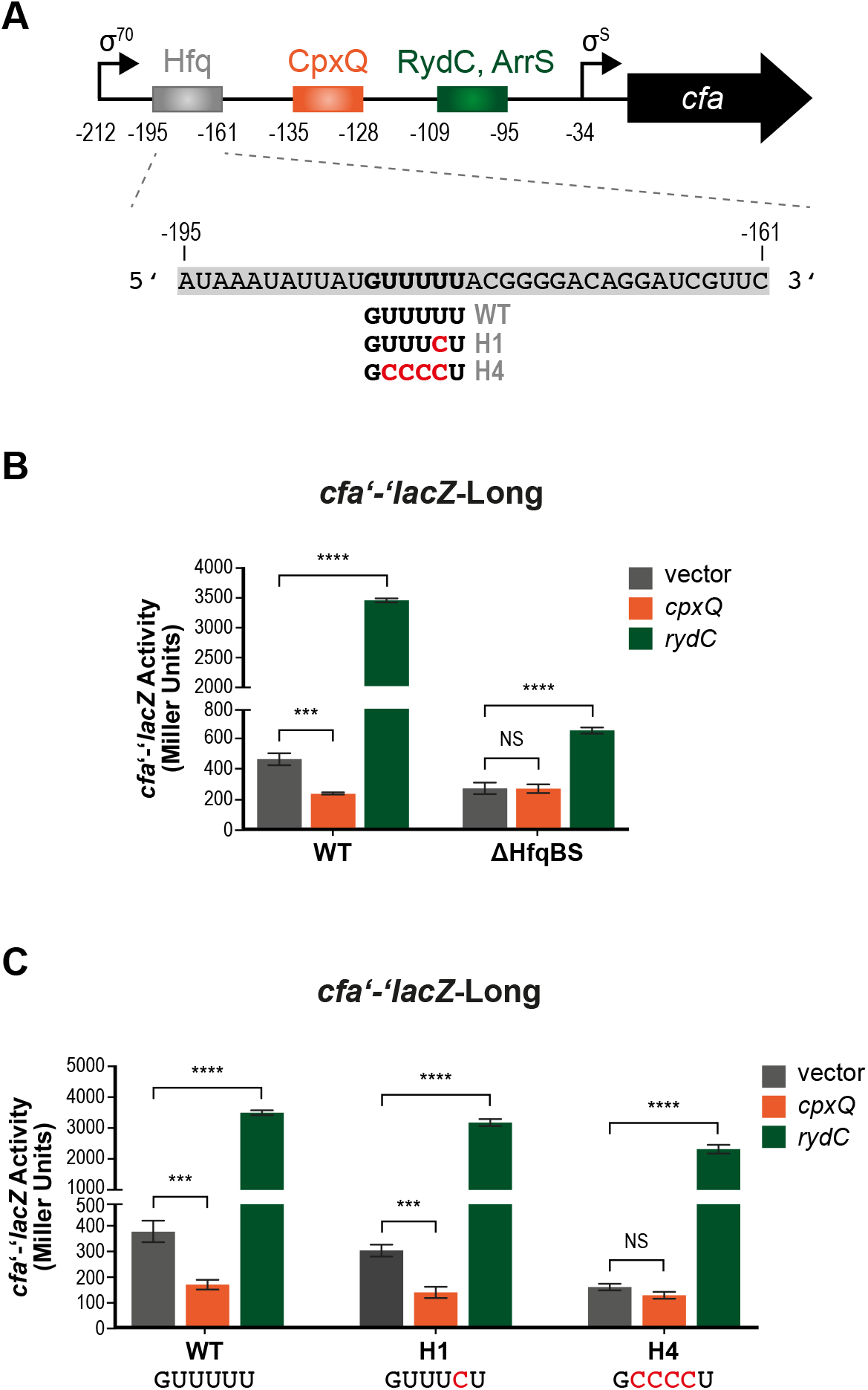
Mutating a putative Hfq binding site alters *cfa* mRNA regulation by CpxQ and RydC. (A) 5’ UTR of *cfa* gene with a putative Hfq binding site and sRNA base pairing sites indicated. (B) The first 50-nt of the 212-nt-5’ UTR (−212 to −163) which includes the putative Hfq binding site, was deleted in *cfa’-’lacZ*-Long fusion (called *cfa’-’lacZ*-ΔHfqBS). Regulation of *cfa’-’lacZ*-Long or *cfa’-’lacZ*-ΔHfqBS by CpxQ and RydC was tested as described in Fig. 1B. (C) Mutations of the Hfq binding site on *cfa* mRNA (shown in A) were made in the *cfa’-’lacZ*-Long fusion background (called *cfa’-’lacZ*-Long H1 and *cfa’-’lacZ*-Long H4). Regulation of WT and mutant fusions by RydC and CpxQ was tested as described in Figure 1B.

To better understand the importance of the −212 to −163 region and Hfq binding site for sRNA-mediated regulation of *cfa* mRNA, we conducted mutational analysis. CLIP-seq captured a U to C crosslinking-induced mutation for *Salmonella* Hfq-cfa mRNA interaction as indicated in the “H1” mutant in Fig. 5A (Holmqvist et al., 2016). The *E. coli* fusion with the single U to C mutation, *cfa’-’lacZ*-Long H1, had a similar level of basal activity to wild-type, and was regulated normally by CpxQ and RydC (Fig. 5C). The sequences of *Salmonella* and *E. coli cfa* mRNAs are not identical in this region (Fig. EV4). Notably, there is a run of U residues in this region in *E. coli* but not in *Salmonella.* To test whether the additional U residues in *E. coli cfa* mRNA contribute to regulation by sRNAs, we also made the “H4” mutant with a total of four U to C substitutions (Fig. 5A). Basal activity of the *cfa’-’lacZ*-Long H4 fusion was reduced compared to the wild-type and H1 fusions (Fig. 5C). Moreover, CpxQ no longer regulated the *cfa’-’lacZ*-Long H4 fusion, but regulation by RydC remained at near wild-type levels (Fig. 5C). This result suggests that Hfq binding at this upstream site may be particularly important for CpxQ-mediated repression and less important for RydC-mediated activation of *cfa.*

### Post-transcriptional regulation of *cfa* mRNA by sRNAs changes membrane lipid composition

We next wanted to assess the consequences of multiple sRNAs regulating cfa expression with regard to the physiology of the bacterium. CFA synthase forms CFAs by transferring a methylene group from S-adenosyl methionine to the double bond of an UFA of a mature phospholipid that is already incorporated in the membrane. Specifically, palmitoleic acid (C16:1) is converted into methylene-hexadecanoic acid (C17CFA) and vaccenic acid (C18:1) is converted into methylene-octadecanoic acid (C19CFA). To test how post-transcriptional regulation of *cfa* by sRNAs alters the proportion of CFAs in membrane fatty acids, we conducted gas chromatography on membrane lipids isolated from strains where each sRNA was ectopically expressed (Fig. 6A, Table S1). The FA composition of all strains was similar except for CFA and UFA content (Fig. 6A, Table S1). In strains carrying *P_lac_-cfa,* we detected reduced levels of 16:1 and 18:1 UFAs and ~6-fold increase in levels of C17CFA, compared to the strain carrying the vector control. Similarly, ArrS-producing cells had ~5-fold higher levels of C17CFA and reduced 16:1 and 18:1 UFAs compared to vector control. RydC-producing cells likewise had reduced levels of 16:1 and 18:1 UFAs and ~8-fold increased levels of C17CFA compared to the control strain. CpxQ-producing strains had modestly reduced C17CFA (~2-fold reduced compared to vector control). These data show that regulation of *cfa* mRNA translation by each sRNA is correlated with changes in membrane CFA content, implying that the regulation by sRNAs could contribute to meaningful changes in cell membrane structure and function.

**Figure 6:**
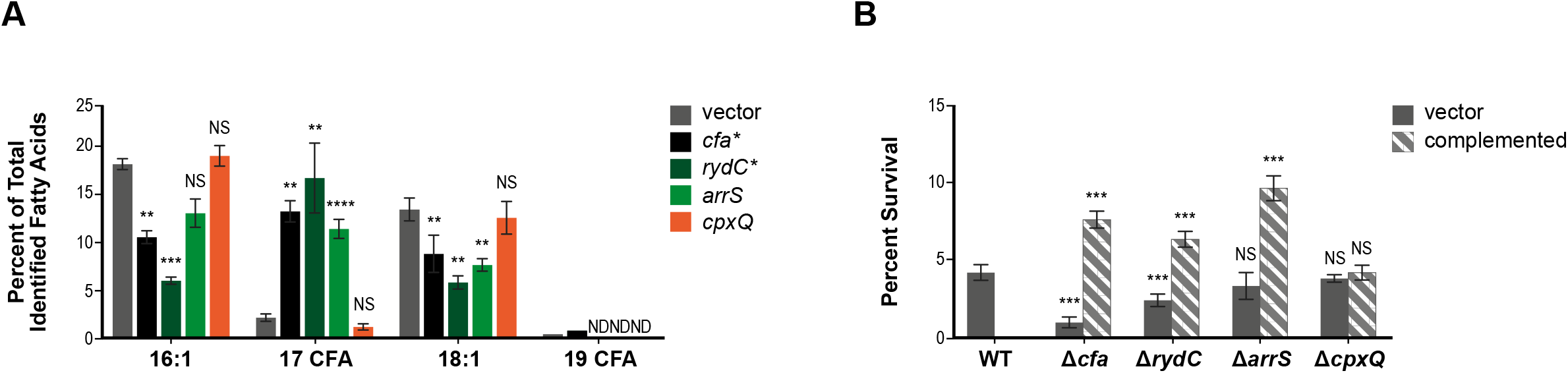
Expression of sRNAs can alter fatty acid composition and survival to acid shock. (A) Relative qualification of fatty acids in *E. coli* in response to ectopic expression of vector control, *cfa, rydC, arrS,* or *cpxQ.* Fatty acids are presented as a percent of total identified fatty acids. Error represents average ± standard deviation, n=3 or *n=2. ND: not detected. The error bars are standard deviations, and the statistical significance was determined using the Student’s *t*-test. \**P* < 0.05, \*\**P* < 0.005, *\*\*\*P* < 0.0005, \*\*\*\**P* < 0.0001, ns is for not significant. The Student’s *t*-test was performed comparing the amount of each fatty acid in response to expression of the sRNA to fatty acid amount in vector control. (B) Survival during an acid challenge. When strains reached an OD_600_ of 0.2, cultures were diluted into LB pH 3 and incubated for 60 minutes. Survival was determined by the ratio of CFUs on the LB plates after acid shock to CFUs on the LB plate before acid shock. Error bars represent average ± standard deviation of 3 technical replicates. The error bars are standard deviations, and the statistical significance was determined using the Student’s *t*-test. \**P* < 0.05, \*\**P* < 0.005, \*\*\**P* < 0.0005, *\*\*\*\*p* < 0.0001, ns is for not significant. The Student’s *t*-test was performed comparing acid survival of each mutant to the survival of vector control.

### Role of sRNAs in surviving acid shock

Because CFAs have been implicated in resistance to acidic pH (Brown et al., 1997, Chang & Cronan, 1999), we next investigated whether sRNAs promote survival after acid shock. Survival of wild-type, *Δcfa, ΔrydC, ΔarrS, ΔgadF* and *ΔcpxQ* strains and complemented mutants was measured after rapid shift of cultures from pH 7 to pH 3. Survival was determined by the ratio of CFUs recovered after acid shock compared to CFUs before acid shock (Fig. 6B). Compared to the wild-type strain, the *ΔrydC,* and *Δcfa* strains had lower survival rates after the acid shock. The *Δcfa* mutant had the lowest survival rate of all the strains tested *(Δcfa* 1.0 ± 0.3 % survival compared to WT 4.3 ± 0.5 %). The *ΔrydC* strain had a survival rate intermediate between that of *Δcfa* and wild-type *(ΔrydC,* 2.5 ± 0.4 %) suggesting that RydC might be important for activating *cfa* or other targets to promote resistance to acid shock. The *ΔarrS,* and *ΔcpxQ* strains showed the same survival rates as wild-type, implying that these two sRNAs do not play an important role in regulating *cfa* or other targets to promote acid shock resistance under these conditions.

Complementation of the *cfa, rydC* and *arrS* mutations by expression of the corresponding gene from a plasmid resulted in enhanced survival of acid shock compared to the wild-type strain (Fig. 6B, “striped” bars). Enhanced acid resistance for ArrS-producing strains was observed previously and was attributed to ArrS positively regulating *gadE* expression (Aiso et al., 2014). Our data suggest that ArrS-dependent activation of *cfa* might also contribute to enhanced acid resistance of ArrS-producing cells. The complemented *cpxQ* strain showed survival rates comparable to the wild-type strain (Fig. 6B). We have shown that ArrS and RydC both activate *cfa* post-transcriptionally and increase the CFA content in cell membranes but only deletion of *rydC* renders otherwise wild-type cells more susceptible to acid shock. These results suggest that at least RydC-mediated activation of *cfa* translation promotes cell survival during an acid shock.

### Small RNA-dependent regulation of *cfa* at acidic pH

To further investigate sRNA-dependent regulation during acid stress, we examined sRNA-dependent regulation of *cfa* translational fusions at neutral and acidic pH. Cells carrying *cfa’-’lacZ*-Long (Fig. 7), *cfa’-’lacZ*-Short or *cfa’-’lacZ*-ΔsRNABS (Fig. EV5) were grown at either pH 7 or pH 5 and assayed for β-galactosidase activity. For *cfa’-’lacZ*-Long, activity was higher at pH 5 compared to pH 7 (Fig. 7). Activity of both *cfa’-’lacZ*-Short and *cfa’-’lacZ*-ΔsRNABS was similar at both pH 5 and pH 7 (Fig. EV5). These observations indicate that activation of *cfa* mRNA in response to acidic pH occurs post-transcriptionally, and the loss of regulation of a fusion lacking the sRNA binding sites suggested that one or more sRNAs could be responsible for regulation under these conditions.

**Figure 7:**
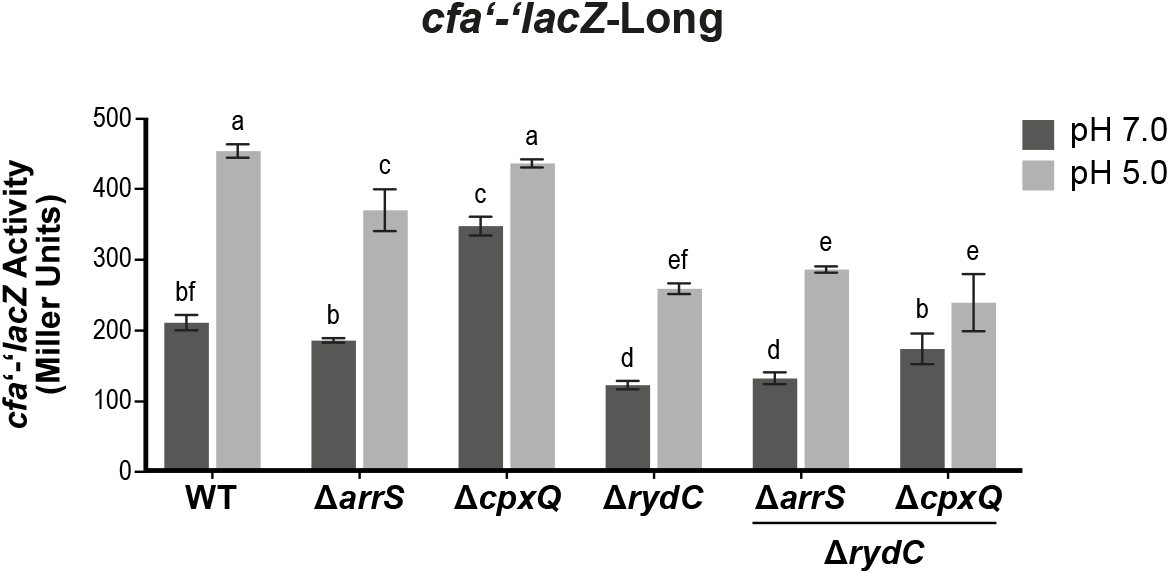
*cfa* translation is induced at acidic pH. Cells carrying *cfa’-’lacZ*-Long in either WT background, a background where one sRNA is deleted, or a background where *rydC* and one other sRNA are deleted were grown in medium at pH 7 then subcultured into medium either pH 7 (pH 7.0) or pH 5 (pH 5.0). Samples were harvested 120 minutes later and assayed for β-galactosidase activity of the reporter fusion. Error bars represent standard deviation for three biological replicates. A one-way ANOVA with post hoc Tukey’s test was performed; the same letter indicates no significant differences between fusion activities (*P* ≤ 0.05).

To further probe the effects of sRNA regulation of *cfa* mRNA at acidic pH, the experiment was performed in wild-type and sRNA mutant backgrounds (Fig. 7). Activity of *cfa’-’lacZ*-Short was not affected by changes in pH or by deletion of any of the sRNAs (Fig. EV5). For *cfa’-’lacZ*-Long, wild-type and *ΔarrS* strains all had similar activities at pH 7 and pH 5, with activity at pH 5 higher than at pH 7 (Fig. 7). The *ΔcpxQ* mutant compared to wild-type had higher *cfa’-’lacZ* activity at pH 7 (Fig. 7). Since ectopic expression of *cpxQ* repressed *cfa* translation (Fig. 1B), the increased levels of *cfa’-’lacZ* activity in the *ΔcpxQ* mutant suggests that at neutral pH, CpxQ is produced at sufficient levels to repress *cfa.* In contrast, at pH 5, the *ΔcpxQ* mutant and wild-type had similar levels of *cfa’-’lacZ*-Long activity (Fig. 7). The *ΔrydC* mutant had lower *cfa’-’lacZ* activity at both pH 7 and 5 compared to wild-type, suggesting that RydC may be produced at sufficient levels at both neutral and acidic pH to have an activating effect on *cfa* under both conditions. We have shown that ectopic production of ArrS or RydC activates *cfa* and increases the CFA content in cell membranes, but only deletion of *rydC* affects *cfa’-’lacZ-Long* activity at pH 5. This observation fits with our previous result that deletion of *rydC* renders otherwise wild-type (*cfa^+^*) cells more susceptible to acid shock (Fig. 6B).

To further investigate the interplay of the sRNAs on *cfa* mRNA translation at different pH values, we deleted *rydC* in combination with each of the other sRNAs in the *cfa’-’lacZ*-Long strain (Fig. 7) and measured β-galactosidase activity of the reporter fusions at pH 5 and 7. Deletion of *arrS* in the *ΔrydC* background had no effect on *cfa’-’lacZ-Long* activity compared to the *ΔrydC* parent (Fig. 7), suggesting that ArrS does not play a role in regulation of *cfa* translation under these conditions. Compared with the *ΔrydC* parent, the *ΔcpxQ ΔrydC* strain had slightly higher *cfa’-’lacZ*-Long activity at pH 7 (Fig. 7). In contrast, at pH 5, the *ΔrydC* parent and *ΔcpxQ ΔrydC* strain had similar levels of *cfa’-’lacZ*-Long activity (Fig. 7). Altogether, the data are consistent with the hypothesis that RydC exerts a positive effect on *cfa* mRNA at both neutral and acidic pH under our growth conditions. In contrast, CpxQ has a mild repressive effect on *cfa* mRNA only at neutral pH.

### RpoS is not responsible for observed differences in *cfa* mRNA regulation

A previous study suggested that transcription from the σ^s^-dependent *cfa* promoter was increased when cells were grown at pH 5 compared to pH 7 (Chang & Cronan, 1999). Our *cfa’-’lacZ*-Long reporter fusion contains the region encompassing the σ^s^-dependent promoter. To determine if the increased *cfa’-’lacZ*-Long activity we observed during mild acid stress was dependent on RpoS, we measured *cfa’-’lacZ*-Long activity at pH 5 and 7 in a *ΔrpoS* background (Fig. 8A). Deletion of *rpoS* had no effect on *cfa’-’lacZ*-Long activity at pH 5 or 7 and we observed the same increase in activity in response to pH 5 in both wild-type and *ΔrpoS* backgrounds (Fig. 8A), indicating that RpoS does not impact the observed activity of the *cfa’-’lacZ*-Long fusion.

**Figure 8:**
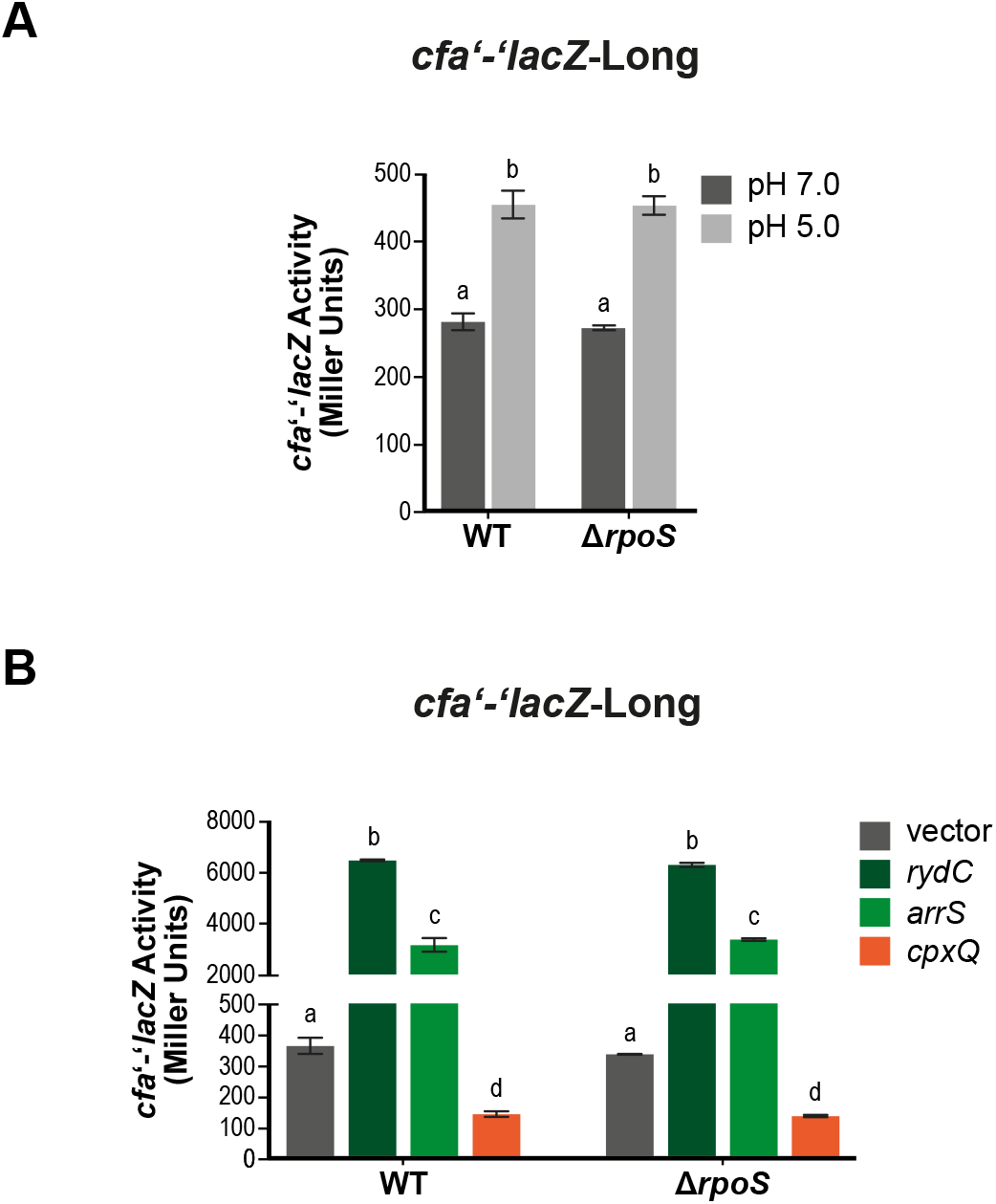
Deletion of *rpoS* does not affect cfa’-’/acZ-Long activity. (A) Cells carrying *cfa’-’lacZ*-Long in a WT or *ΔrpoS* background were grown and *cfa* translation in response to acid was assayed as described in Figure 7. (B) *cfa’-’lacZ*-Long in a WT or *ΔrpoS* background carrying an empty, P_lac_-arrS, P_lac_-cpxQ, P_lac_-gadF, or P_lac_-rydC plasmid were grown as described in Figure 1B.

We showed that ectopic production of each sRNA, RydC, ArrS, and CpxQ can regulate *cfa’-’lacZ*-Long (Fig. 1B). While the experiment above suggests that pH-dependent regulatory effects are not mediated by RpoS, we wanted to further test whether sRNAs could be acting indirectly on *cfa* via post-transcriptional regulation of *rpoS.* We ectopically expressed RydC, ArrS, or CpxQ in wild-type and *ΔrpoS *cfa’-’lacZ**-Long strains and did not observe any differences in sRNA-dependent regulation between wild-type and *ΔrpoS* (Fig. 8B), indicating that the sRNA-dependent regulation is not mediated indirectly through RpoS. These data further support the idea that the regulation of *cfa* by these sRNAs is direct.

## Discussion

Microbial membranes are the first line of defense against environmental stress as well as the site of many metabolic processes. As such, maintenance of membrane integrity and homeostasis is key to cell survival. Small RNAs (sRNAs) are important post-transcriptional regulators that play key roles in the response to environmental stress. We are interested in how sRNAs can contribute to membrane modification in response to stressors, particularly how sRNAs can lead to altered phospholipid composition. In the present work, we investigate the roles of multiple sRNAs in post-transcriptional regulation of *cfa* mRNA, encoding a key enzyme used to alter lipid composition of bacterial membranes. We show that *cfa* can be post-transcriptionally regulated by four sRNAs, all of which act on the same long isoform of *cfa* mRNA that results from transcription from the constitutive σ^70^-dependent promoter (Fig. 9A). RydC and ArrS post-transcriptionally activate *cfa,* while the sRNA CpxQ post-transcriptionally represses *cfa* (Fig. 1B). RydC and ArrS bind at a site that overlaps an RNase E cleavage site, which may prevent RNase E cleavage, increase *cfa* mRNA stability and allow for increased *cfa* translation. How CpxQ represses *cfa* translation is still unknown. Based on the location of the CpxQ-cfa mRNA base pairing interaction, it is unlikely that the repression mechanism is similar to those that have already been characterized for CpxQ. We hypothesized that CpxQ could be working through Hfq or RNase E (Fig. 9A). Our mutational analyses indicate that both the RNase E recognition site that is adjacent to the site for base pairing by activating sRNAs (Fig. 4) and an Hfq binding site (Fig. 5) affect *cfa* mRNA stability and are necessary for CpxQ repression. We propose that CpxQ base pairing with *cfa* mRNA makes the RNase E cleavage site more susceptible to degradation. One possibility is that CpxQ:cfa base pairing induces a structural change that makes this RNase E site more susceptible to cleavage. CpxQ may also increase degradation at this RNase E site by directing Hfq to one binding site at the expense of another (Fig 9A). RydC requires Hfq to stabilize *cfa* mRNA (Fröhlich et al., 2013) but does not require the identified Hfq binding site, indicating that there may be another Hfq binding site in the 5’ UTR of *cfa.* The sRNAs may determine the location of Hfq binding which in turn may modulate the accessibility of the RNase E cleavage site. Further studies on RNase E cleavage of *cfa* mRNA in the presence of CpxQ or Hfq CLIP-seq in *E. coli,* will give more insight into how Hfq and RNase E affect CpxQ repression of *cfa* mRNA.

**Figure 9:**
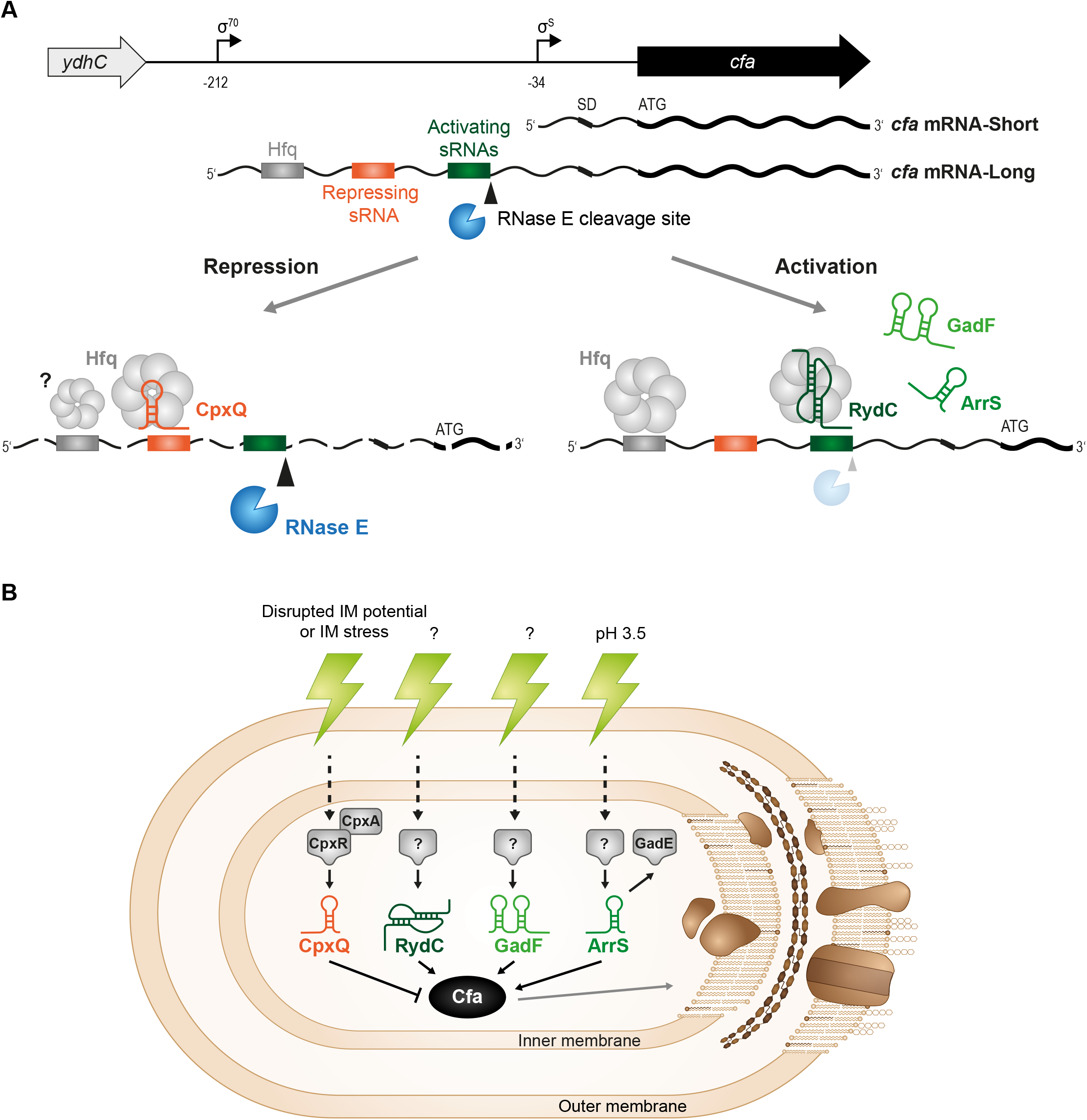
Models of sRNAs regulation of *cfa* mRNA. (A) Working model for the role of Hfq, RNAse E and sRNAs in regulation of *cfa* mRNA. There are multiple RNase E cleavage sites that have been mapped in *Salmonella.* Activating sRNAs, especially RydC, require Hfq for efficient pairing and inhibition of RNAse E cleavage at one site that may be the primary cleavage site, and the pairing stabilizes *cfa* mRNA. CpxQ may have the opposite effect – somehow pairing of CpxQ (facilitated by Hfq), stimulates RNAse E-dependent cleavage especially at the site RydC is known to protect. (B) Regulation of *cfa* mRNA by these sRNAs provides a sigmaS-independent mechanism for control of cfa, and thus a way to modulate lipid composition of the bacterial cell envelope in response to a variety of different stresses. CpxQ is part of the Cpx regulon, which responds to signals like disrupted IM potential or IM stress. The conditions that stimulate RydC production are unknown. ArrS and GadF are linked to acid stress, and CFAs have previously been implicated in acid resistance. Modulating the composition of cyclopropanated FA in response to these stresses could alter membrane properties like fluidity and permeability and allow the cell to adapt to the stress.

Regulation of *cfa* mRNA by multiple sRNAs suggests that under some condition(s) these sRNAs influence membrane lipid composition and membrane integrity to promote fitness or stress resistance. Towards elucidating the physiological roles of RydC, ArrS, and CpxQ in the regulation of membrane composition, we investigated the role of each sRNA in acid stress, a condition where CFA synthase is known to play a role (Brown et al., 1997, Chang & Cronan, 1999). We determined that *cfa* translation was higher when cells were exposed to pH 5 compared to pH 7 (Fig. 7). When *rydC* was deleted, *cfa* translation could not reach maximum wild-type levels at pH 5 and *ΔrydC* mutants were more susceptible to acid shock compared to wild-type, giving us the first phenotype associated with RydC (Figs. 6B, 7). However, it is not currently known if activation of *cfa* at acidic pH is the major physiological role of RydC, as the conditions and signals stimulating RydC production are not yet known.

While ectopic expression of *arrS* increased *cfa* translation and promoted higher levels of acid resistance (Figs. 1B, 6B), deletion of *arrS* did not affect *cfa* translation or acid shock survival (Figs. 6B, 7), indicating that ArrS may not play a role in acid stress under the conditions tested. ArrS has been characterized as an activator of the transcriptional activator GadE and ectopic expression of ArrS increases acid resistance though its activation of *gadE* but a deletion phenotype has not been reported (Aiso et al., 2014, Aiso et al., 2011). ArrS may activate *cfa* under another condition where CFAs in the membrane contribute to stress resistance (Fig. 9B).

Expression of GadF only slightly regulated *cfa* reporter fusions (Fig. 1B). GadF is relatively uncharacterized but was shown to activate *acrB* 12-fold compared to the reported 2-fold activation of *cfa* (Melamed et al., 2016). (Note, in our hands, the regulation of *cfa* by GadF is even more modest, at ~1.3-fold activation (Fig. 1B).) AcrB is an efflux pump that uses proton motive force to pump compounds, such as antibiotics, organic solvents, and detergents, out of the cell (Anes, McCusker et al., 2015). GadF regulation of *cfa* mRNA may be relevant under these conditions, as CFA membrane content has been shown to promote resistance to organic solvents (Kanno, Katayama et al., 2013). Understanding the regulation of GadF itself would provide insight into the importance of its regulation of *cfa* mRNA, however, it is not currently known how or in response to what stimuli *gadE* mRNA (from which GadF is derived) is processed (Fig. 9B).

CpxQ is the only sRNA of the four we have characterized that represses *cfa* (Fig. 1B). Deletion of *cpxQ* resulted in increased *cfa* expression at pH 7 compared to wild-type but did not affect *cfa* at pH 5 (Fig. 7). The physiological consequences of CpxQ-mediated repression of *cfa* remain unclear. Responding to acid stress is likely not a relevant physiological role for CpxQ since deletion of *cpxQ* did not affect acid shock survival (Fig. 6B). However, our observation that CpxQ represses *cfa* translation at neutral pH combined with what is known about CpxQ may provide some insight. CpxQ is part of the Cpx two-component system that responds to cell envelope stress (Chao & Vogel, 2016, Grabowicz et al., 2016). Both the Cpx two-component system and the processing of CpxQ are induced when cell membrane potential is dissipated (Chao & Vogel, 2016). The translation of the two targets of CpxQ, an inner membrane sodium-proton antiporter NhaB and the periplasmic chaperone Skp, is repressed, possibly to minimize membrane stress and maintain the proton motive force (PMF) across the inner membrane (Chao & Vogel, 2016, Grabowicz et al., 2016, Grabowicz & Silhavy, 2017).

CpxQ regulation of *cfa* might also help to maintain inner membrane PMF, but the effect of CFAs on maintenance of membrane potential is currently unknown. A previous study showed that a lack of CFAs in cellular membranes increases H+ permeability but decreases H+ extrusion (Shabala & Ross, 2008). Furthermore, the Cpx response is widely thought to be important for the regulation of energy production and substrate transport across the inner membrane (Raivio, 2014) and fatty acid composition is known to affect inner membrane transporter activity and protein function (Lee, 2004, Tveld, Driessen et al., 1993) but how CFAs affect these functions has not been studied. However, there is a linear correlation between enzyme activity and membrane fluidity (Tveld et al., 1993) and, *in vitro,* CFAs are known to increase membrane fluidity by disrupting lipid packing (Poger & Mark, 2015).

Another possibility that has not been explored is the effect of the Cfa protein and mechanism of catalysis on inner membrane integrity. During catalysis, one subunit of the Cfa dimer binds to the membrane while the other subunit extracts and modifies unsaturated lipids from the membrane (Hari, Grant et al., 2018). Given that the Cpx response regulates events at the inner membrane and predominantly inhibits the production of inner membrane localized protein complexes, Cfa’s tight association and interaction inside the inner membrane may trigger the Cpx response. Lastly, PMF is the cellular energy source and the conversion of UFAs to CFAs by Cfa is an energetically expensive reaction (Wang & Cronan, 1994) so CpxQ may repress *cfa* translation when PMF is dissipated simply to reduce cellular ATP consumption. Clearly defining the impact of Cfa on PMF and determining under what conditions CpxQ differentially regulates its regulon would elucidate the physiological function of CpxQ regulation of *cfa* translation.

The UFA to CFA conversion by CFA synthase is energetically expensive, so it is perhaps not surprising that CFA synthase production is tightly controlled at the transcriptional level (Kim et al., 2005, Wang & Cronan, 1994) and at the post-transcriptional level by multiple sRNAs. Regulation of *cfa* mRNA by multiple sRNAs may have many advantages. There are a few examples of multiple sRNAs acting on the same target. Post-transcriptional control of the important transcription factor RpoS (σ^s^) by multiple sRNAs to mediate responses to diverse stress conditions is well documented (Battesti, Majdalani et al., 2011). The *rpoS* gene is post-transcriptionally regulated by at least four sRNAs in response to different conditions. sRNA regulation of *rpoS* highlights the potential importance of regulation of the same mRNA target by multiple sRNAs: synthesis of each sRNA is induced by a different stress which allows RpoS in turn to properly respond to these various stress signals. We propose that expression of each sRNA is induced by a different environmental condition that either requires more Cfa, in the case of RydC, ArrS, and GadF, or less Cfa in the case of CpxQ (Fig. 9B). It is known that transcription of *arrS* is induced by acid stress and that CpxQ is produced in response to cell membrane potential dissipation, but how that relates to their regulation of *cfa* is not understood. The regulation of *rydC* itself has not been elucidated, and it is not currently known how *gadE* is processed or in response to what stimuli. Uncovering the stimuli that promote synthesis of ArrS, CpxQ, GadF, and RydC would provide more insight into how and why *cfa* mRNA is regulated by multiple sRNAs.

Extensive previous work has shown that sRNAs play an integral role in changing membrane protein composition in response to changing environments (Grabowicz & Silhavy, 2017, Guillier & Gottesman, 2006, Holmqvist & Wagner, 2017, Vogel & Papenfort, 2006) but how sRNAs influence membrane fatty acid composition is not well studied. Here, we build on the first example of an sRNA altering fatty acid composition (Fröhlich et al., 2013) by showing regulation of *cfa* by four sRNAs. Currently, the sRNAs studied here are the only examples of sRNAs altering the fatty acid composition of membranes, which can consequently alter the stability and permeability of the membrane.

## Materials and Methods

### Strain and plasmid construction

Strains and plasmids used in this study are listed in Table EV2. All strains used in this study are derivatives of *E. coli* K12 DJ624 (D. Jin, National Cancer Institute). Oligonucleotide primers used in this study are listed in Table EV3. Integrated DNA Technologies synthesized all primers. *ΔrpoS* and *Δcfa* mutations were made via P1 *vir* transduction from the Keio collection (Baba, Ara et al., 2006). All other chromosomal mutations were made using λ Red recombination (Datsenko & Wanner, 2000, Yu, Ellis et al., 2000) and marked alleles were moved between strains by P1 *vir* transduction (Miller & H, 1972). Kanamycin markers were removed using pCP20 (Datsenko & Wanner, 2000).

The *lacZ* translational reporter fusions were constructed using λ Red homologous recombination into strain PM1205 and counter-selection against *sacB,* as previously described (Mandin & Gottesman, 2009). Transcription of the fusion is under control of the *P_BAD_* promoter. The fusion contains the *cfa* 5’ UTR fragment of interest and the first 36 codons of the *cfa* open reading frame. *cfa’-’lacZ*-Long, *cfa’-’lacZ*-Short, *cfa’-’lacZ*-ΔsRNABS, and *cfa’-’lacZ-*ΔHfqBS were constructed by PCR amplifying the *cfa* 5’ UTR fragment of interest from DJ624 using a fusion-specific forward primer containing flanking homology to P_BAD_ and the reverse primer O-AK24R, which contains flanking homology to *lacZ* and anneals in the *cfa* coding region (Table EV3). *cfa’-’lacZ*-Long and *cfa’-’lacZ*-Short were constructed by A.M. King (King et al., 2019). *cfa’-’lacZ*-Long H1 and H4 were constructed by using a forward primer containing the desired point mutations and flanking homology to P_BAD_ and reverse primer O-AK24R to PCR amplify the *cfa* 5’ UTR fragment of interest from DJ624 (Table EV3). *cfa’-’lacZ*-LongM1 and *cfa’-’lacZ-*LongM2 were made using Gibson Assembly (Gibson, Young et al., 2009). pBR322 was PCR amplified using primers with 5’ homologies to the *cfa* 5’ UTR containing the desired mutations. Gibson reaction was performed using NEBuilder HiFi DNA Assembly Master Mix (New England Biolabs) according to the manufacture’s protocol. The Gibson product was transformed into XL10 competent cells and the plasmid was purified. The mutated *cfa* 5’ UTR was PCR amplified from this plasmid using the same primers used to create *cfa’-’lacZ-Long.* The plasmid used to make *cfa’-’lacZ*-LongM1, which contains the G101C point mutation, was saved as pCB1 for further strain construction. The RNase E recognition site mutant fusions *cfa’-’lacZ*-LongG1 and G2 were made by mutating pCB1 using QuikChange mutagenesis (Agilent Technologies) and oligos that restore the G101C mutation to the WT and add the additional desired mutations. The plasmid used to make *cfa’-’lacZ-LongG2* was saved as pCB2. *cfa’-’lacZ*-LongG3 and G4 were made by mutating pCB2 using QuikChange mutagenesis (Agilent Technologies) and oligos that add the desired mutations. The mutated *cfa* 5’ UTR was PCR amplified from each plasmid using the same primers used to create *cfa’-’lacZ*-Long. All PCR products were recombined into PM1205 using λ Red homologous recombination and counter-selection against *sacB* as previously described (Mandin & Gottesman, 2009). All fusions were verified using DNA sequencing.

Plasmids containing WT sRNAs, mutant sRNAs, and Cfa under the control of the P_LlacO_ promoter were constructed by PCR amplifying each sRNA from *E. coli* DJ624 chromosomal DNA using oligonucleotides containing HindIII and BamHI restriction sites (Table EV3). The primers used to create P_LlacO_-arrS *M1* and P_LlacO_-rydC *M1* contained the desired point mutations. PCR products and vector pBRCS12 (Wadler & Vanderpool, 2009) were digested with HindIII and BamHI (New England Biolabs) restriction endonucleases. Digestion products were ligated using DNA Ligase (New England Biolabs) and the plasmids were transformed into XL10 competent cells and the plasmid was purified. Pı__lacO_-cpxQ *M2* was created using QuikChange mutagenesis (Agilent Technologies) of PLlacO*-cpxQ* using primers in Table EV3. All plasmids were confirmed by DNA sequencing.

### Media and growth conditions

Bacteria were cultured in LB broth medium or on LB agar plates at 37°C, unless stated otherwise. When necessary, media were supplemented with antibiotics at following concentrations: 100μgml^-1^ ampicillin (Amp) or 25μgml^-1^ kanamycin (Kan). Isopropyl β-D-1-thiogalactopyranoside (IPTG) was used at 0.1 mM (final concentration) for induction of expression from P_LlacO-1_ promoter.

### β-Galactosidase assays

#### Ectopic expression of sRNAs

Bacterial strains harboring translational *lacZ* reporter fusions carrying the sRNA plasmid were cultured overnight in Terrific Broth (TB) with Amp and 0.002% L-arabinose then subcultured 1:100 to fresh TB medium containing Amp and 0.002% L-arabinose. Strains were grown at 37°C with shaking to early exponential phase then 0.1 mM IPTG was added to induce sRNA expression. Samples were harvested 60 minutes later and β-galactosidase assays were then performed as previously described (Miller & H, 1972).

#### Acid

Bacterial strains harboring translational *lacZ* reporter fusions were cultured overnight in TB medium pH 7.0 with 0.002% L-arabinose then subcultured 1:100 to fresh TB medium pH 7.0 containing 0.002% L-arabinose. Strains were grown at 37°C with shaking for 2 hours then strains were subcultured 1:100 again into TB medium with 0.002% L-arabinose at pH 7.0 or pH 5.0. Samples were harvested 120 minutes later and β-galactosidase assays were then performed as previously described (Miller & H, 1972).

### Northern Blot

Bacterial strains were cultured in LB medium to OD_600_~0.4 then plasmids were induced with 0.1 mM IPTG. After 60 minutes, total RNA was extracted by the hot phenol method as described previously (Aiba, Adhya et al., 1981). RNA concentrations were measured spectrophotometrically.15 μg of RNA were denatured for 5 min at 95°C in loading buffer (containing 95% formamide), separated on 8% polyacrylamide urea gel at 100 V for 1 h using 1× Tris-acetate-EDTA (TAE), then transferred to BrightStar™-Plus Positively Charged Nylon Membrane in 0.5× TAE buffer by electroblotting at 50 V for 1 hour at 4°C. RNA was crosslinked to the membrane then membrane was prehybridized for 45 min in ULTRAhyb (Ambion) solution at 42°C. Blots were hybridized overnight with P^32^-end-labeled primer “CpxQNB” then membranes were washed at 42°C twice with 2× SSC/0.1% SDS for 8 minutes then twice with 0.1× SSC/0.1% SDS for 15 minutes. Signal was detected using film. Membranes were stripped in boiling 0.1% SDS for 10 minutes then re-probed overnight P^32^-end-labeled primer “5S.” Ladder was prepared using the Decade Markers System from ThermoFisher, according to manufacture’s instructions.

### Analysis of fatty acids

Cells were cultured overnight in LB containing Amp and subcultured 1:100 to fresh LB containing Amp. Cells were grown to an OD_600_ of 0.1 then plasmids were induced with 0.1M IPTG for one hour. Cells were then spun down and the cell pellet was washed twice with MilliQ water. Fatty acids were extracted according to a previously described protocol, method 3.1 (Dionisi, Golay et al., 1999). Pentadecanoic acid methyl ester was used as an internal standard. The bacterial sample was spiked with 500 μg of the internal standard, pentadecanoic acid, and then the sample was transesterified with 2 mL of 0.5M sodium methoxide for 1 min at room temperature. Fatty acid methyl esters (FAMEs) were then extracted using 2 mL of hexane. The solution was centrifugation at 2000 rpm for 5 min and the organic upper phase was removed and dried under nitrogen. FAMEs were treated with trimethylsilyldiazomethane in methanol to ensure complete methylation. FAMEs were identified on an Agilent 6890N GC with a 5973 MS and a Zebron WAX column (30m × 0.25mm × 0.25μm).

### Acid Shock Assay

Strains were grown overnight in LB with Amp then subcultured 1:100 into LB with Amp at pH 7.0 and grown at 37°C with shaking until OD_600_= 0.1. Plasmids were induced with 0.1mM IPTG. When strains reached an OD_600_ of 0.2, cultures were diluted 10x into LB pH 3.0 and incubated at 37°C with no shaking for 60 minutes. Survival was determined by the ratio of CFUs on the LB plates after acid shock to CFUs on the LB plate before acid shock.

### *In vitro* RNA synthesis and structure probing

DNA templates carrying a T7 promoter sequence for *in vitro* RNA synthesis were generated by PCR from *E. coli* gDNA *(cfa* mRNA: KFO-0702/KFO-0703; RydC KFO-0704/KFO-0705; ArrS KFO-0708/KFO-0709; GadF KFO-0710/KFO-0711; CpxQ KFO-07406/KFO-0707). 200 ng of template DNA were transcribed using the AmpliScirbe T7 Flash Transcription Kit (Epicentre) following the manufacturer’s recommendations; size and integrity of the transcripts were verified on a denaturing 6% PAA gel.

RNA structure probing was carried out as described previously (Fröhlich et al., 2013) with few modifications. In brief, 0.4 pmol 5’ end-labelled *cfa* mRNA were mixed with 0.4 pmol *E. coli* Hfq protein (provided by K Bandyra and BF Luisi, University of Cambridge) or Hfq dilution buffer (1X structure buffer, 1% (v/v) glycerol, 0.1% (v/v) Triton X-100) in the presence of 1X structure buffer (0.01 M Tris pH 7, 0.1 M KCl, 0.01 M MgCl_2_) and 1 μg yeast RNA, and samples were incubated at 37°C for 10min. Subsequently, unlabelled sRNA (4 pmol) or water was added, and reactions were kept at 37°C for an additional 10min. Samples were treated with RNase T1 (0.1 U; Ambion, #AM2283) for 2 min or with lead(II) acetate (final concentration: 5mM; Sigma #316512) for 1.5 min, respectively.

### Primer extension

Primer extension experiments were performed as previously described (Fröhlich et al., 2013). A 5’ end-labelled primer (KFO-0920) specific to *gfp* was used for reverse transcription of the samples, as well as for the preparation of a template specific ladder (prepared using the USB Cycle Sequencing Kit; affymetrix USB #78500; template amplification pZE-Cat/KFO-0920 on pKF206).

## Acknowledgements

We would like to express our gratitude to Dr. Sandy Pernitzsch from Scigraphix for creating figures. We greatly appreciate discussions and experimental advice from Dr.

Joel Belasco and Dr. Jamie Richards. Thank you to Dr. Alexander Ulanov from the Roy J. Carver Biotechnology Center for analyzing the fatty acids. We are grateful to Vanderpool lab members, both past and present, for strains, plasmids, discussions, and advice. We appreciated Dr. James Slauch and the members of his lab for valuable discussions. This work was supported by the National Institutes of Health (R01 GM092830) to C.K.V. and supported C.M.B. K.S.F. acknowledges funding by the LMU Mentoring program of the LMU Faculty of Biology.

## Author contributions

C.M.B., K.S.F., and C.K.V. designed research; C.M.B. and K.S.F. performed research; C.M.B., K.S.F., and C.K.V. analyzed data; and C.M.B., K.S.F., and C.K.V. wrote the paper.

## Expanded View Figures

**Figure EV1:**
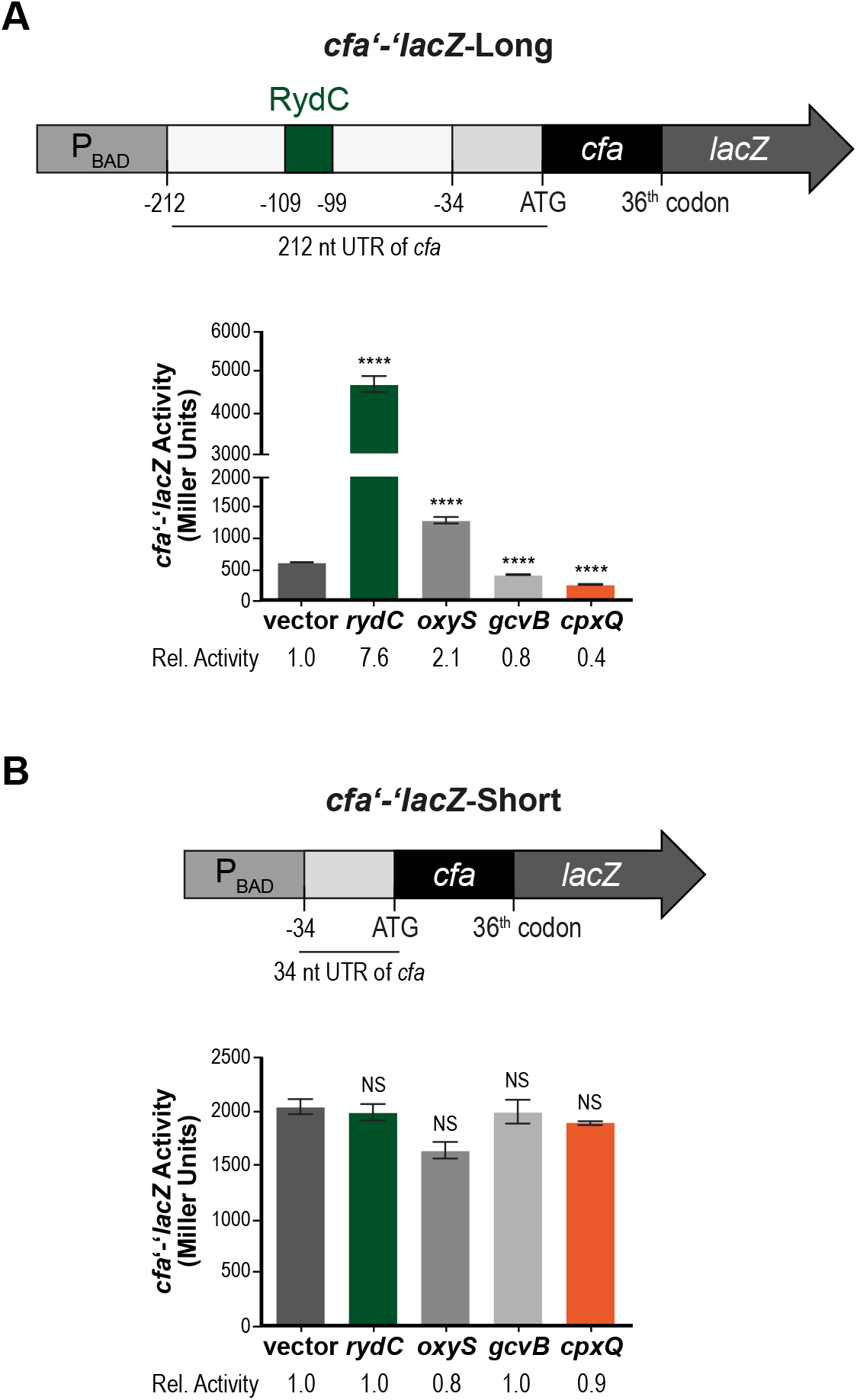
Testing of additional sRNAs for regulation of *cfa* expression. (A) The *cfa’-’lacZ*-Long fusion contains the entire 212-nt 5’ UTR, including the RydC binding site (indicated by green square labeled “RydC”). Regulation of *cfa’-’lacZ*-Long by RydC, OxyS, GcvB, and CpxQ was determined as described in Fig. 1B. (B) The *cfa’-’lacZ*-Short fusion contains only proximal the σ^s^-dependent promoter. Regulation of this fusion by RydC, OxyS, GcvB, and CpxQ was determined as described in Fig. 1B.

**Figure EV2:**
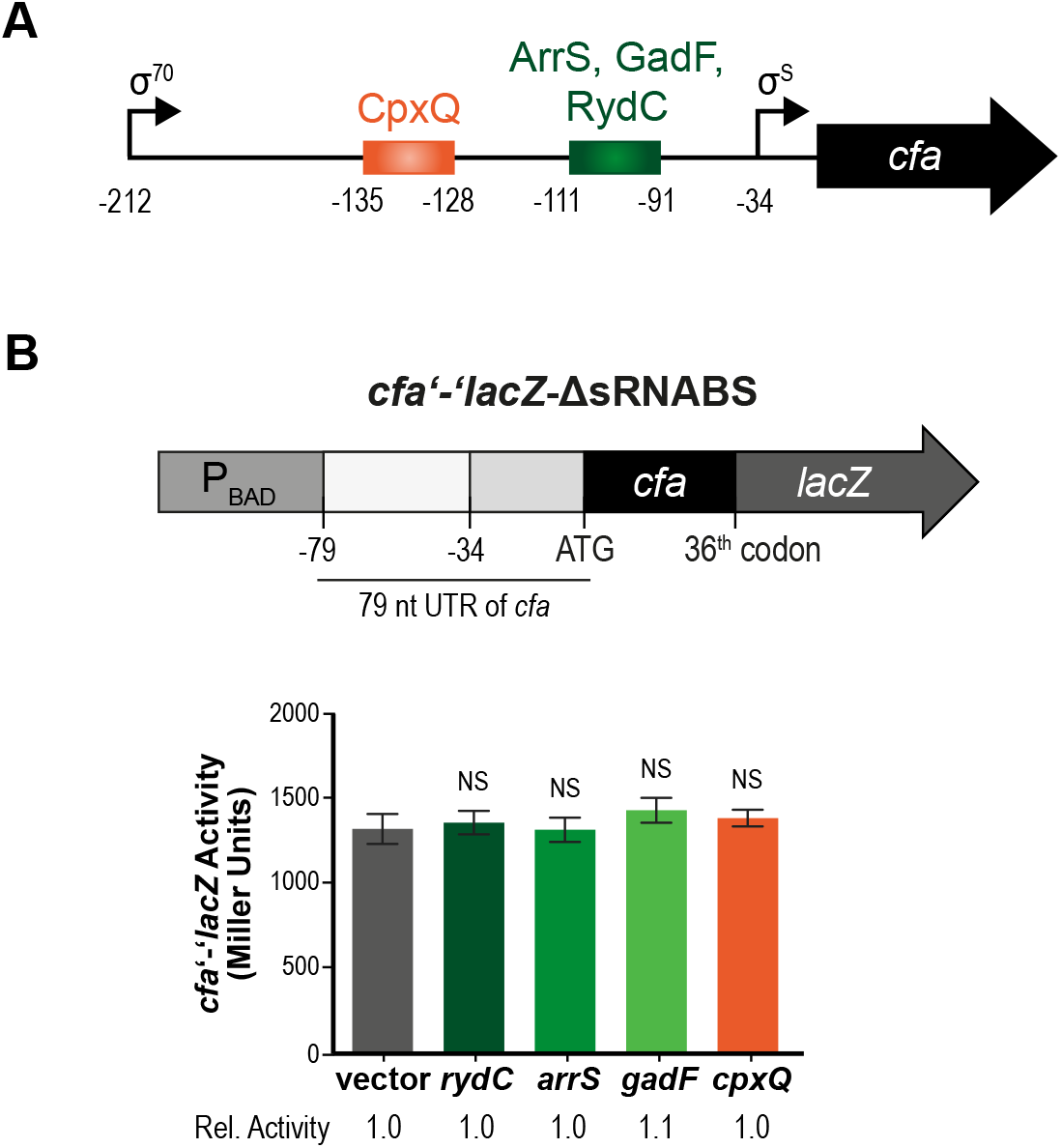
Deletion of the putative sRNA binding site removes regulation of *cfa* translation by ArrS, CpxQ, GadF, and RydC. (A) 5’ UTR of *cfa* gene. Arrows mark transcriptional start sites and sRNA binding sites are indicated with labeled boxes. (B) A *cfa* translational fusion to *lacZ* (cfa’-’lacZ-ΔsRNABS) that begins immediately downstream of the predicted sRNA binding site was constructed. This fusion contains the proximal σ^s^-dependent promoter and 79-nt upstream of this promoter. (C) Regulation of *cfa’-’lacZ*-ΔsRNABS by ArrS, CpxQ, GadF, or RydC was tested as described in Figure 1B.

**Figure EV3:**
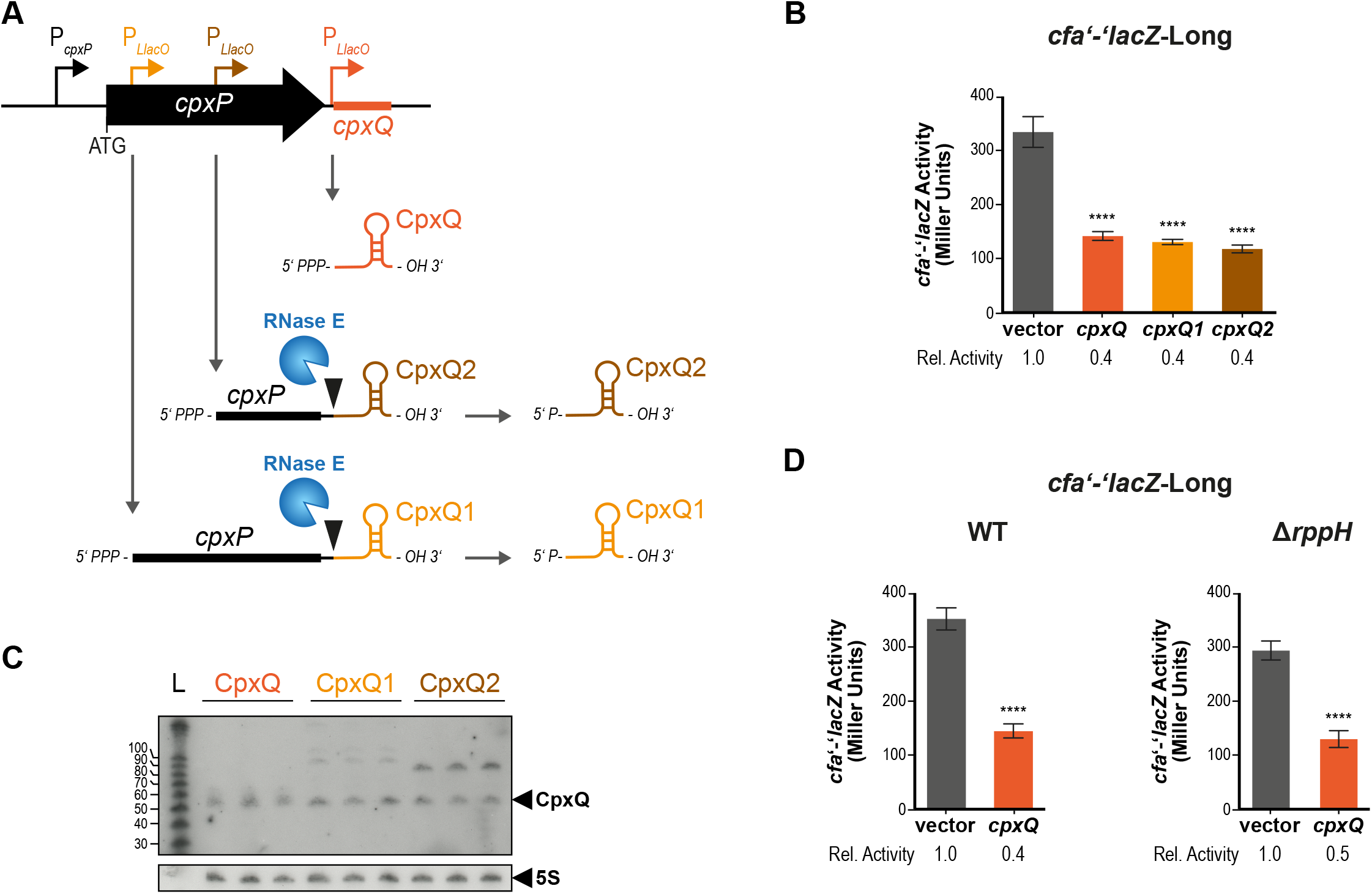
CpxQ does not require a 5’ monophosphate to repress *cfa.* (A) Three CpxQ plasmids were constructed: CpxQ encodes CpxQ directly from the CpxQ start site, CpxQ1 starts right after the “AUG” of CpxP and CpxQ2 starts 30-nt upstream of the RNase E cleavage site in the 3’ UTR of CpxP. Plasmids were designed so that CpxQ1 and CpxQ2 would be cleaved by RNase E to yield 5’ P CpxQ. (B) Regulation of *cfa’-’lacZ*-Long was tested for each sRNA plasmid as described in Fig. 1B. (C) Expression levels of CpxQ from each plasmid as described in (A) were determined by northern blot analysis of total RNA samples. 5S served as loading control. (D) Regulation by CpxQ was tested for *cfa’-’lacZ*-Long in a WT or *ΔrppH* background as described in Fig. 1B.

**Figure EV4:**
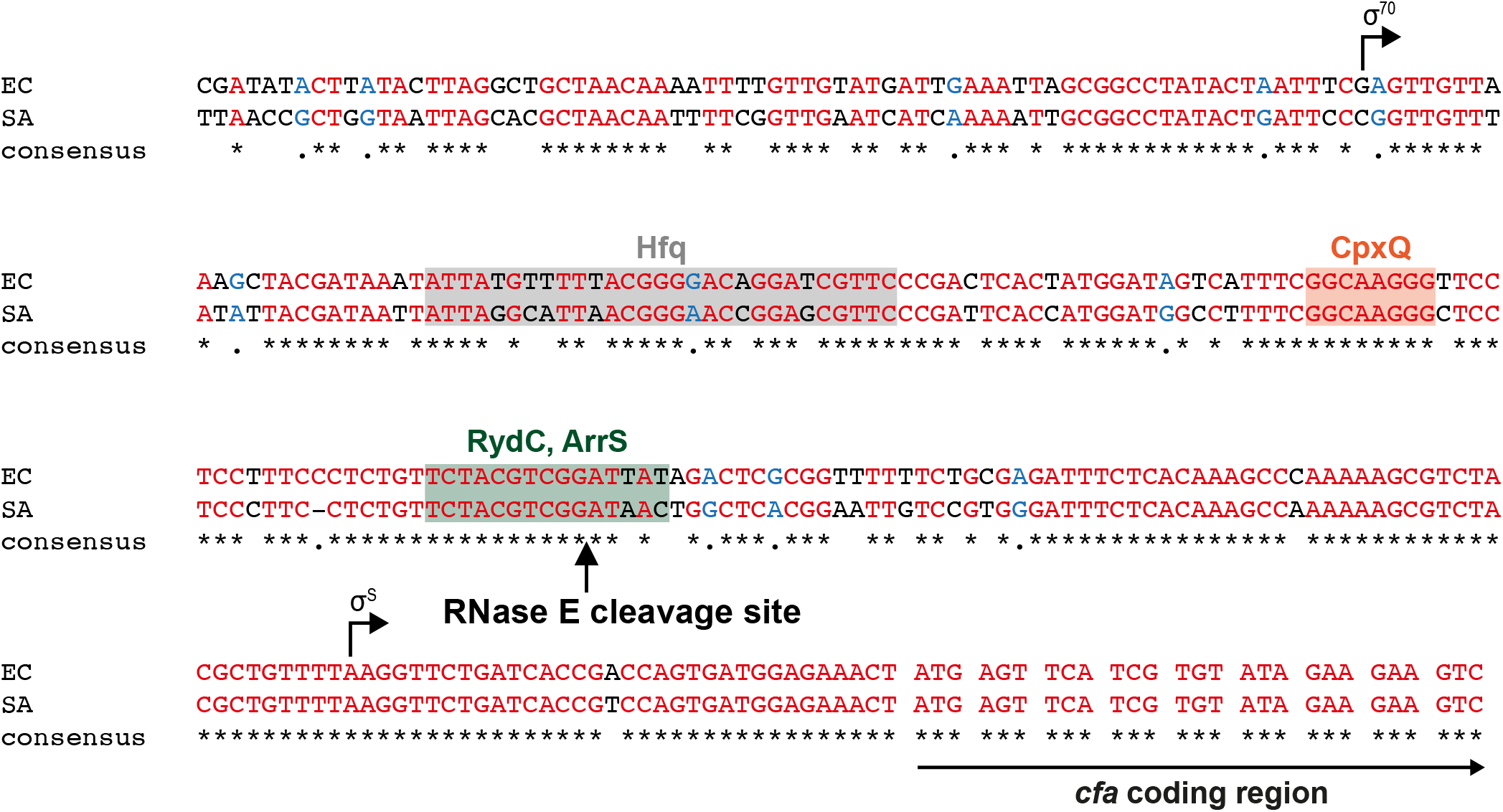
The *cfa* 5’ UTR differs between *E. coli* and *Salmonella*. Sequence alignment of the promoter and 5’ UTR of *cfa* in *E. coli* and *Salmonella.* Gray box: Hfq binding site; Orange box: CpxQ binding site. Green box: Activating sRNA binding site. Black arrow: RNase E cleavage site.

**Figure EV5:**
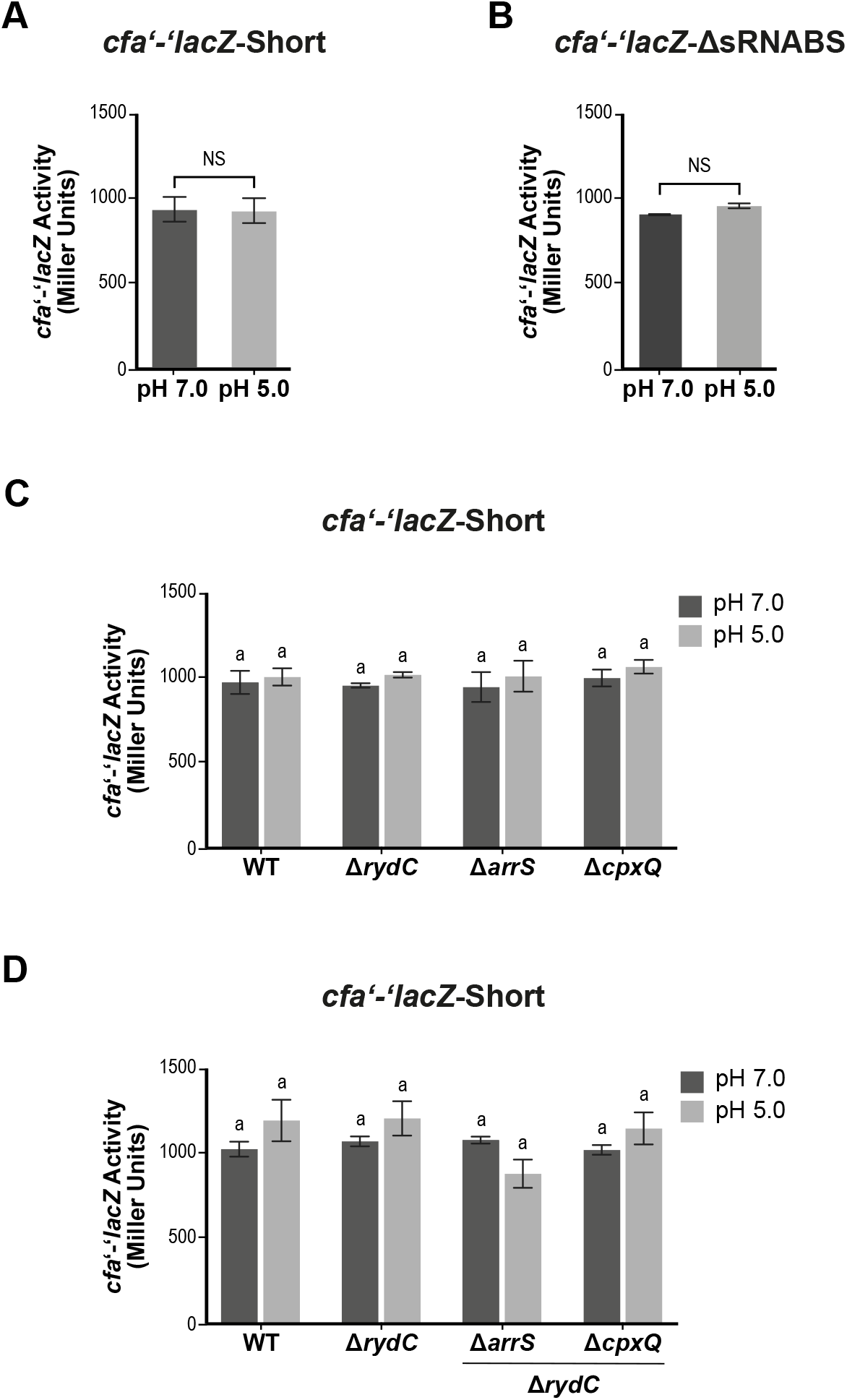
Deletion of sRNAs do not affect *cfa’-’lacZ-Short* activity. (A) Cells carrying *cfa’-’lacZ*-Short were grown and *cfa* translation in response to acid was assayed as described in Figure 7. (B) Cells carrying *cfa’-’lacZ*-ΔsRNABS were grown and *cfa* translation in response to acid was assayed as described in Figure 7. (C) Cells carrying *cfa’-’lacZ*-Short in either WT background or a background where one sRNA is deleted were grown and *cfa* translation in response to acid was assayed as described in Figure 7. (D) Cells carrying *cfa’-’lacZ*-Short (in either WT background or a background where *rydC* and one other sRNA are deleted) were grown as described in Figure 7. *ΔrydC* single mutant is included for reference. Error bars represent standard deviation for three biological replicates. A one-way ANOVA with post hoc Tukey’s test was performed; the same letter indicates no significant differences between fusion activities (*P* ≤ 0.05).

## Supplemental Tables/ Excel files

**Table S1:**
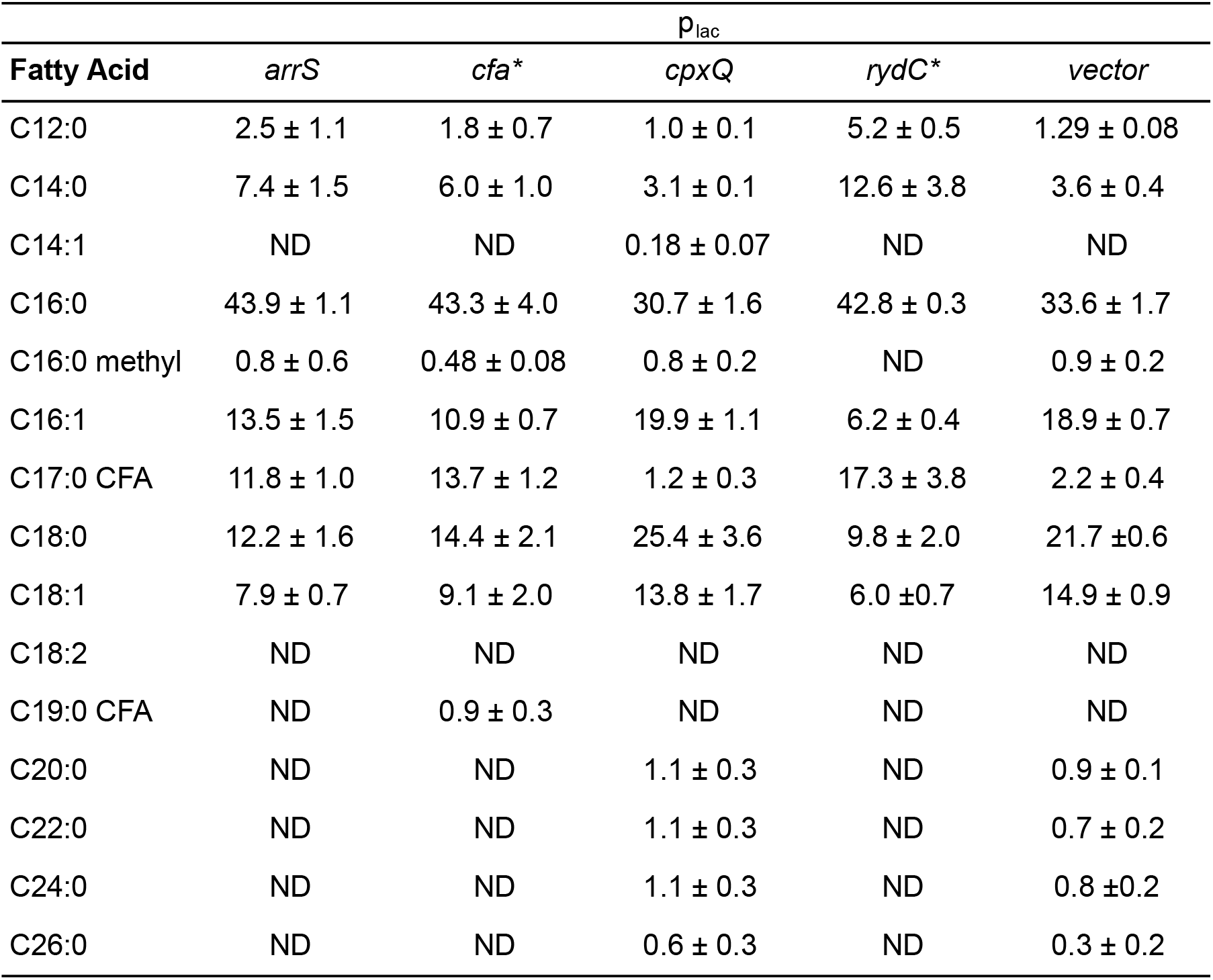
Expression of sRNAs can alter fatty acid composition. Relative qualification of fatty acids in *E. coli* in response to ectopic expression of *arrS, cfa, cpxQ, rydC* or vector control. Fatty acids are presented as a percent of total identified fatty acids. Error represents average ± standard deviation, n=3 or *n=2. ND: not detected

| Fatty Acid | P<sub>lac</sub> |  |  |  |  |
| --- | --- | --- | --- | --- | --- |
|  | *arrS* | *cfa* * | *cpxQ* | *rydC* * | *vector* |
| C12:0 | 2.5 ± 1.1 | 1.8 ± 0.7 | 1.0 ± 0.1 | 5.2 ± 0.5 | 1.29 ± 0.08 |
| C14:0 | 7.4 ± 1.5 | 6.0 ± 1.0 | 3.1 ± 0.1 | 12.6 ± 3.8 | 3.6 ± 0.4 |
| C14:1 | ND | ND | 0.18 ± 0.07 | ND | ND |
| C16:0 | 43.9 ± 1.1 | 43.3 ± 4.0 | 30.7 ± 1.6 | 42.8 ± 0.3 | 33.6 ± 1.7 |
| C16:0 methyl | 0.8 ± 0.6 | 0.48 ± 0.08 | 0.8 ± 0.2 | ND | 0.9 ± 0.2 |
| C16:1 | 13.5 ± 1.5 | 10.9 ± 0.7 | 19.9 ± 1.1 | 6.2 ± 0.4 | 18.9 ± 0.7 |
| C17:0 CFA | 11.8 ± 1.0 | 13.7 ± 1.2 | 1.2 ± 0.3 | 17.3 ± 3.8 | 2.2 ± 0.4 |
| C18:0 | 12.2 ± 1.6 | 14.4 ± 2.1 | 25.4 ± 3.6 | 9.8 ± 2.0 | 21.7 ± 0.6 |
| C18:1 | 7.9 ± 0.7 | 9.1 ± 2.0 | 13.8 ± 1.7 | 6.0 ± 0.7 | 14.9 ± 0.9 |
| C18:2 | ND | ND | ND | ND | ND |
| C19:0 CFA | ND | 0.9 ± 0.3 | ND | ND | ND |
| C20:0 | ND | ND | 1.1 ± 0.3 | ND | 0.9 ± 0.1 |
| C22:0 | ND | ND | 1.1 ± 0.3 | ND | 0.7 ± 0.2 |
| C24:0 | ND | ND | 1.1 ± 0.3 | ND | 0.8 ± 0.2 |
| C26:0 | ND | ND | 0.6 ± 0.3 | ND | 0.3 ± 0.2 |

**Table S2:** Plasmids and Strains used in this study

**Table S3:** Oligos used in this study

